# Thrombospondin-1 inhibits alternative complement pathway activation in vasculitis synergistically to factor H

**DOI:** 10.1101/2024.07.31.606000

**Authors:** Swagata Konwar, Sophie Schroda, Jessika Kleindienst, Eva L. Decker, Martin Pohl, Barbara Zieger, Jens Panse, Hong Wang, Robert Grosse, Manuel Rogg, Christoph Schell, Sabine Vidal, Xiaobo Liu, Christian Gorzelanny, Todor Tschongov, Karsten Häffner

**Affiliations:** Department of Internal Medicine IV (Nephrology), Medical Center, Faculty of Medicine, University of Freiburg, Freiburg, Germany; Department of General Pediatrics, Adolescent Medicine and Neonatology, Medical Center, Faculty of Medicine, University of Freiburg, Freiburg, Germany; Plant Biotechnology, Faculty of Biology, University of Freiburg, Freiburg, Germany; Division of Pediatric Hematology and Oncology, Department of Pediatrics and Adolescent Medicine, Faculty of Medicine, Medical Center-University of Freiburg, 79098 Freiburg, Germany; Department of Oncology, Hematology, Hemostaseology and Stem Cell Transplantation, University Hospital RWTH Aachen, Aachen, Germany; Center for Integrated Oncology (CIO), Aachen, Bonn, Cologne, Düsseldorf (ABCD) Germany Pauwelsstrasse 30, 52074, Aachen, Germany; Institute of Experimental and Clinical Pharmacology and Toxicology, Medical Faculty, University of Freiburg, 79104 Freiburg, Germany; Centre for Integrative Biological Signalling Studies-CIBSS, University of Freiburg, 79104 Freiburg, Germany; Institute of Surgical Pathology, Medical Center, Faculty of Medicine, University of Freiburg, 79106 Freiburg, Germany; Freiburg Institute for Advanced Studies (FRIAS), University of Freiburg, 79106 Freiburg, Germany; Department of Dermatology and Venereology, University Medical Center Hamburg-Eppendorf, Hamburg, Germany

## Abstract

Complement activation is a relevant driver in the pathomechanisms of vasculitis. The involved proteins in the interaction between endothelia, complement and platelets in these conditions are only partially understood.

Thrombospondin-1 (TSP-1), found in platelet α-granules and released from activated endothelial cells, interacts with factor H (FH) and von Willebrand factor (vWF). However, direct regulatory interaction with the complement cascade has not yet been described.

We could show that TSP-1 is a potent, FH-independent inhibitor of the alternative complement pathway. TSP-1 binds to complement proteins, inhibits cleavage of C3 and C5 and the formation of the membrane attack complex. Complement-regulatory function is validated in blood samples from patients with primary complement defects. Physiological relevance of TSP-1 is demonstrated in ANCA-vasculitis patients by significantly enhanced TSP-1 staining in glomerular lesions and increased complement activity and NETosis following TSP-1 deficiency in an ANCA-vasculitis model.

The newly described complement-inhibiting function of TSP-1 represents an important mechanism in the interaction of endothelia, complement and platelets. In particular, the interplay between released TSP-1, vWF and the complement system locally, especially on surfaces, influences the balance between complement activation and inhibition and may be relevant in various vascular diseases.

## Introduction

Thrombospondins (TSPs) are multifunctional, matricellular glycoproteins consisting of 5 members (TSP-1, TSP-2, TSP-3, TSP-4 and TSP-5) (1). TSP-1 is the first and most prominent member of this protein family. It is the major component of the α-granules in platelets (up to 25%) and can additionally be released from the Weibel-Palade bodies of endothelial cells (2, 3). TSP-1 is also produced and secreted after tissue injury by a variety of cells including endothelial cells, fibroblasts and macrophages (4). TSP-1 is involved in the regulation of clot formation and platelet aggregation (5). Furthermore, TSP-1 stabilizes platelet aggregates by inhibiting vWF-mediated proteolysis by ADAMTS13 (6, 7). TSP-1 supports wound healing through the activation of latent TGF-β and has been involved in a number of fibrotic diseases like diabetic nephropathy, liver fibrosis and multiple myeloma (8). It also has been shown to be a potent inhibitor of the NO signaling cascade through its interaction with the cell surface receptor CD47 (9).

Until now, complement regulatory functions have only been shown for TSP-5 in rheumatoid arthritis, inhibiting the classical pathway (10). Considering the similar structure of all thrombospondins, other members of this family could also possess complement regulatory functions. Especially TSP-1 is likely to interact with the complement system due to its presence in platelets and endothelial cells. In addition, TSP-1 has been shown to co-purify with complement factor H (FH), the most important regulator of the alternative system, isolated from platelets (11). Surface plasmon resonance (SPR) studies have shown that TSP-1 can bind to FH with high affinity, thereby increasing the binding of FH to platelets (12).

Defects in the alternative complement pathway (AP) regulation can predispose to the onset of several severe diseases (13). In atypical hemolytic uremic syndrome (aHUS) complement overactivation on endothelial surfaces, mostly caused by *factor H* mutations, promotes thrombotic microangiopathy and consequently hemolytic anemia, thrombocytopenia and acute and chronic renal failure (14). In paroxysmal nocturnal hemoglobinuria (PNH) acquired mutations in the *phosphatidylinositol glycan class A gene* on hematopoietic stem cells lead to a loss of CD55 and CD59, important negative complement regulators on red cell membranes (15). Consequently these erythrocytes are not sufficiently protected from complement-mediated lysis. While some patients are only affected by mild anemia, a significant amount of untreated patients develop blood-clots that can result in life-threatening thromboembolic events and represent the leading cause of death from this condition (16).

As these direct complement-driven diseases demonstrate, complement activation on surfaces leads to an interplay between the endothelium, complement, platelets, and the coagulation system (17–20). In addition, emerging studies from basic and clinical research show an interaction between these systems even in non-primary complement diseases. For example SARS-CoV-2 hyperinflammation and thrombosis have been linked to complement activation or antineutrophil cytoplasmatic antibody-associated vasculitis (ANCA-vasculitis) (21, 22).

In SARS-CoV-2, the spike protein is able to attach to endothelial surfaces, causing endothelial cell injury and alternative pathway complement activation by competing with factor H for binding to heparan sulfates (23). This leads to endothelial cell activation and complement deposition on microvascular endothelial cells. In addition, large vWF multimers as a platform for complement activation have been shown to contribute to the cytokine storm associated with SARS-CoV-2 (24). Several clinical trials have been conducted investigating whether complement inhibition could prove beneficial in COVID-19 (25, 26).

In ANCA-vasculitis, patients develop autoantibodies against neutrophil antigens that cause inflammation, infiltration of immune cells, and necrosis of small blood vessels (27). Animal models (28) have demonstrated a clear link between vascular inflammation and alternative pathway complement activation, which led to the development and clinical implementation of the C5a receptor inhibitor Avacopan (29).

A deeper understanding of the proteins involved in this interplay between endothelium, complement and platelets could lead to the identification of new pathomechanisms and generate new therapeutic strategies to improve patient health and survival. Considering the relationship of TSP-1 to FH and vWF and its involvement in inflammation, wound healing, and thrombosis, we investigated the ability of TSP-1 to directly interact with the complement system. We examined binding of TSP-1 to alternative pathway complement proteins besides FH and deciphered its complement regulatory functions *in vitro*. In healthy and patients’ blood samples, this complement inhibitory function was verified.

The interaction of TSP-1, vWF and the complement system was investigated in stimulated endothelial cells. In ANCA-vasculitis patients, the relevance of TSP-1 regarding disease activity was examined. These investigations provide evidence for an additional local complement regulatory function of TSP-1 especially in vasculitis, representing one important player in these conditions which opens up new aspects for therapeutic interventions.

## Results

### TSP-1 has factor H-independent complement-regulatory function on the alternative complement pathway

We used recombinant full-length TSP-1 and elucidated its ability to inhibit the alternative complement pathway (AP) by ELISA in normal human serum.

Complement activation was assessed using microtiter plates coated with specific activators of the AP and detecting the amount of membrane attack complexes (MAC) formed in the wells. TSP-1, but not TSP-5, showed pronounced inhibition of the AP at concentrations above 400 nM (Figure 1A). To ensure that the observed experimental effect is not influenced by the histidine tag of recombinant TSP-1, platelet-isolated TSP-1 (p-TSP-1) was also employed in this experiment, yielding comparable results (Figure 1A).

**Figure 1:**
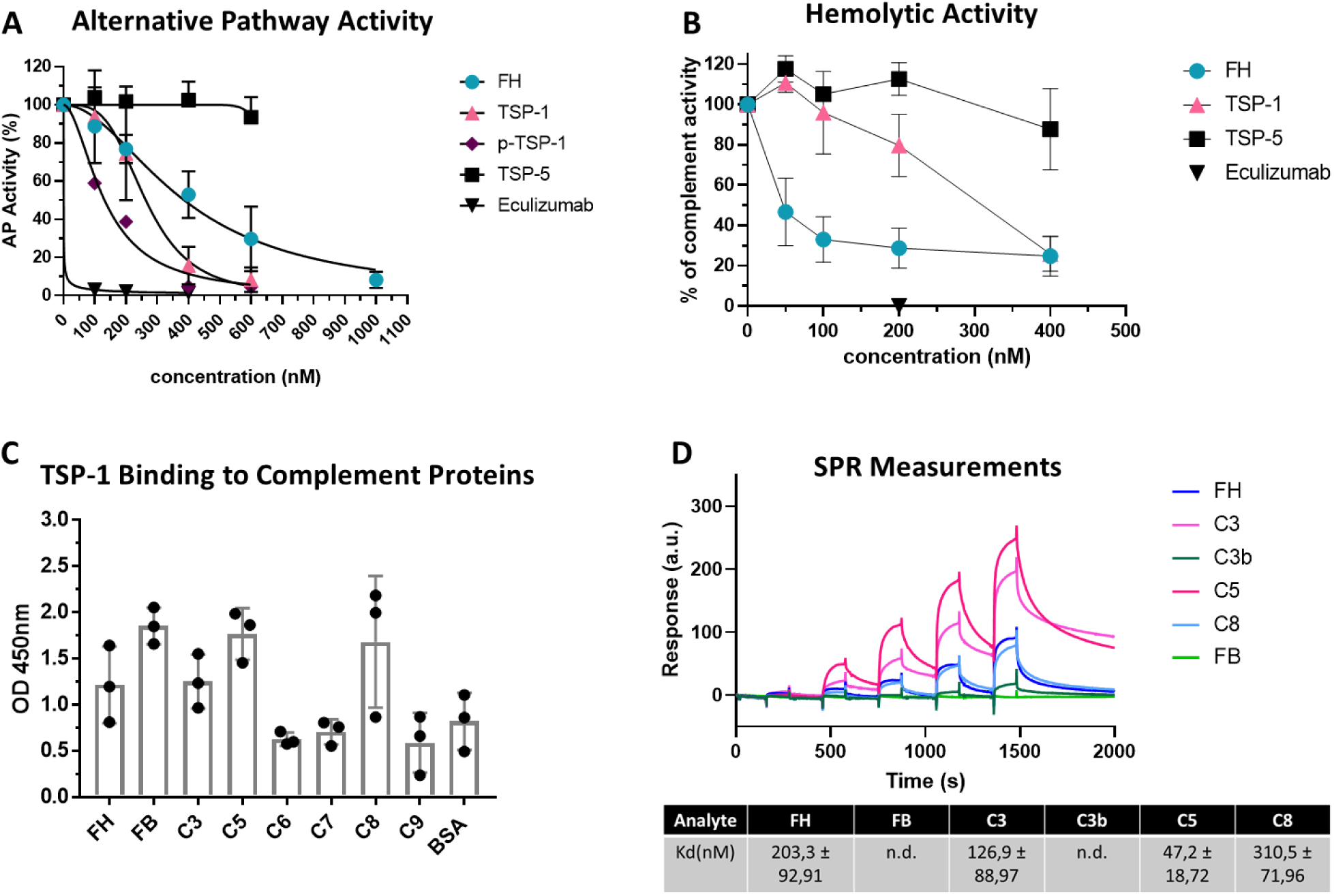
TSP-1 has distinct complement regulatory roles compared to TSP-5, protects sheep erythrocytes from complement mediated hemolysis independent of FH and binds to central complement proteins of the alternative pathway. **(A)** TSP-1 inhibits AP activation in contrast to TSP-5. AP in NHS was activated on LPS coated wells and with increasing concentrations of FH, Eculizumab, TSP-1 or TSP-5. Platelet derived TSP-1 (p-TSP-1) was used as an additional control to exclude artifacts caused by the histidine tag used for purification. **(B)** TSP-1 protects sheep erythrocytes from AP mediated lysis in the absence of FH. Sheep erythrocytes were incubated with FH depleted serum and increasing concentrations of FH, TSP-1 or TSP-5. Results are expressed as means ± SD. AP and hemolytic activity were normalized against untreated control samples. Data was fitted using nonlinear regression. **(C)** TSP-1 binds to central proteins of the alternative pathway. Complement proteins FH, FB, C3, C5, C6, C7, C8, C9 or BSA were coated on microtiter plates and incubated with recombinant TSP-1. Bound TSP-1 was determined using specific antibodies. **(D)** Surface Plasmon Resonance (SPR) Biacore measurements demonstrating the binding of TSP-1 to key proteins of the alternative complement pathway. TSP-1 was immobilized on CM5 chips at a concentration of 0.1 µM. Binding interactions with complement proteins FH, FB, C3, C3b, C5, and C8 were assessed at various concentrations (12.3, 37.03, 111.1, 333.3, 1000 nM). The binding data were fitted using a 1:1 Langmuir binding model to determine on and off rates, which were then used to calculate affinity constants (Kd). The graph depicts a summary of the binding of complement proteins to TSP-1 at increasing concentrations. Average Kd values, calculated from three repeated measurements, are presented in the accompanying table. Data is represented as mean ± SD

The significance of the interaction between TSP-1 and FH was assessed through a hemolytic assay involving sheep erythrocytes and FH-depleted human serum which leads to hemolysis of the erythrocytes due to FH deficiency. TSP-1, in contrast to TSP-5, was able to reduce hemolysis (Figure 1B). The inhibition of hemolysis reached approx. 70 % at concentrations of 400 nM. The fact that TSP-1 could suppress hemolysis in FH-depleted serum implies that TSP-1’s inhibition of the AP is, to some extent, independent of FH.

### TSP-1 binds to central complement proteins of the alternative pathway

In order to explore potential interactions between TSP-1 and the complement system, we investigated the binding of TSP-1 to AP proteins using ELISA. Our findings revealed that TSP-1 bound to FH, FB, C3, C5 and C8 (Figure 1C). To validate these interactions, binding ELISAs were conducted with platelet-isolated TSP-1, eliminating potential binding artifacts introduced by the histidine tag of the protein (supplemental Figure S1). For a comprehensive understanding of binding affinities, we performed SPR Biacore measurements for TSP-1 with FH, FB, C3, C3b, C5, and C8. The strongest binding was observed with C5 (47 nM), followed by C3 (127 nM), FH (203 nM) and C8 (311 nM) (Figure 1D). Although binding to FB and C3b was noted, the affinity was insufficient to determine Kd values within our tested concentration ranges. Given these interactions with several AP proteins, we subsequently employed several *in vitro* assays to specifically assess TSP-1-dependent complement inhibition.

### TSP-1 inhibits factor D-mediated cleavage of factor B

Complement activation involves the cleavage of factor B (FB) by factor D (FD) in the presence of C3b to build up the C3 convertase. Therefore, we investigated whether TSP-1 could inhibit the cleavage of FB. C3b was incubated with FB, FD and increasing amounts of TSP-1. Without TSP-1, almost all FB (93 kDa) underwent cleavage by FD into Ba (33 kDa) and Bb (60 kDa). However, as the concentration of TSP-1 increased, a pronounced decrease in the cleavage products of FB was observed (Figure 2A). This finding indicates that TSP-1 interacts with and protects FB from FD-mediated cleavage.

**Figure 2:**
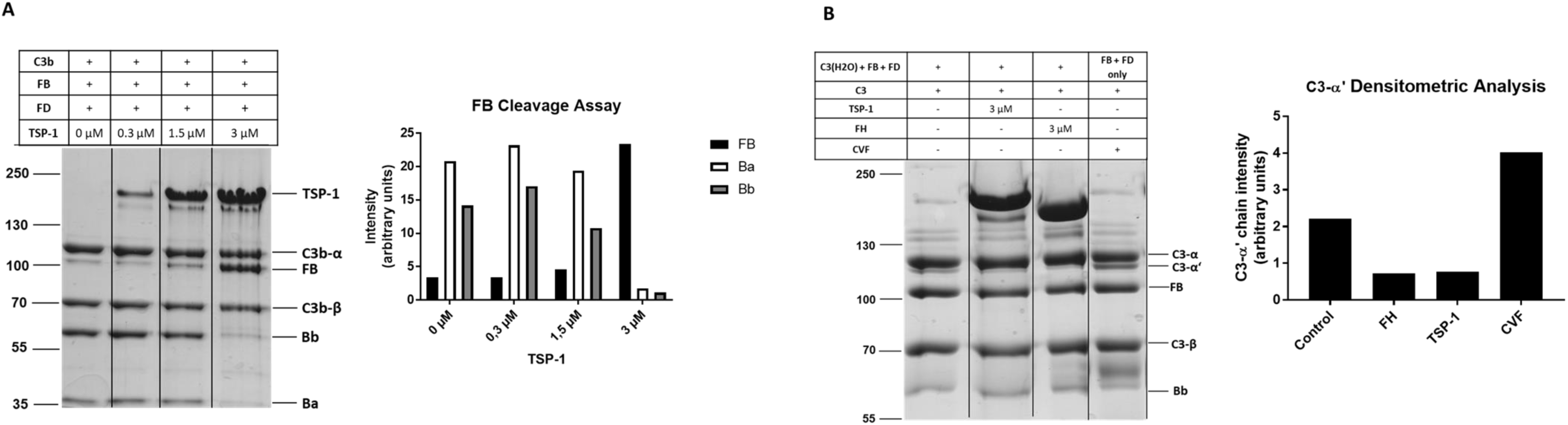
TSP-1 modulates complement at the C3 level of the complement cascade. **(A)** TSP-1 protects FB from cleavage by FD. FB was incubated with FD, C3b, and varying concentrations of TSP-1. Subsequently, FB and its cleavage products were visualized by coomassie staining. Graph on the right illustrates band intensity of FB and its cleavage products. **(B)** TSP-1 inhibits cleavage of C3 by the AP C3 convertase. C3 convertase was generated by incubating C3(H2O) with FB and FD, followed by the addition of C3 in the presence or absence of TSP-1 or FH. As a positive control, CVF C3 convertase (CVF + FB + FD) was utilized. The graph on the right depicts the band intensity of C3α‘ chain. Results are shown as means.

### TSP-1 prevents the cleavage of C3 by the C3 convertase

To determine the ability of TSP-1 to influence C3 cleavage, C3 convertase was generated by incubating C3(H_2_O) with FB and FD. C3 was then added to the reaction in the presence or absence of TSP-1, FH or cobra venom factor (CVF) and the amount of cleaved C3 products was visualized. C3 is a 185 kDa protein consisting of a C3α chain (110 kDa) and a C3β chain (75 kDa) linked by two disulfide bonds. The C3 convertase cleaves the C3α chain into C3a (9 kDa) and the C3-α’ chain (101 kDa). A pronounced C3-α‘ chain was observed in the presence of CVF and in the control sample without any inhibitor (Figure 2B). TSP-1 or FH reduced the amount of generated C3bα’ chain compared to the control sample, indicating that both protein inhibited the cleavage of C3.

### TSP-1 prevents the cleavage of C5 and the formation of the MAC

To assess the functional relevance of the interaction between TSP-1 and C5, we employed a CVF convertase assay. It involved the incubation of C5 in the presence or absence of FH, Eculizumab, TSP-1 or MFHR1 (a recombinant FH-FHR1 fusion protein known for inhibiting C5 cleavage and MAC formation) (30). The quantification of produced C5a was carried out using ELISA. Unlike FH, TSP-1 was able to reduce C5a generation, similarly to Eculizumab and MFHR1. This suggests that TSP-1 is able to protect C5 from cleavage into C5a and C5b (Figure 3A). The formation of the MAC marks the endpoint of the complement cascade. Due to the interaction of TSP-1 with C8, we investigated whether TSP-1 could directly influence MAC formation beyond inhibiting C5 cleavage. The MAC forms when C5b is present on surfaces along with C6-C9. To explore TSP-1’s potential inhibition of direct MAC formation, TSP-1 was mixed with C7, C8 and C9 and then added to sheep erythrocytes preincubated with C5b. In contrast to Eculizumab, TSP-1 demonstrated a robust inhibition of MAC formation and lysis of sheep erythrocytes, comparable to MFHR1 as a positive control (Figure 3B) (30).

**Figure 3:**
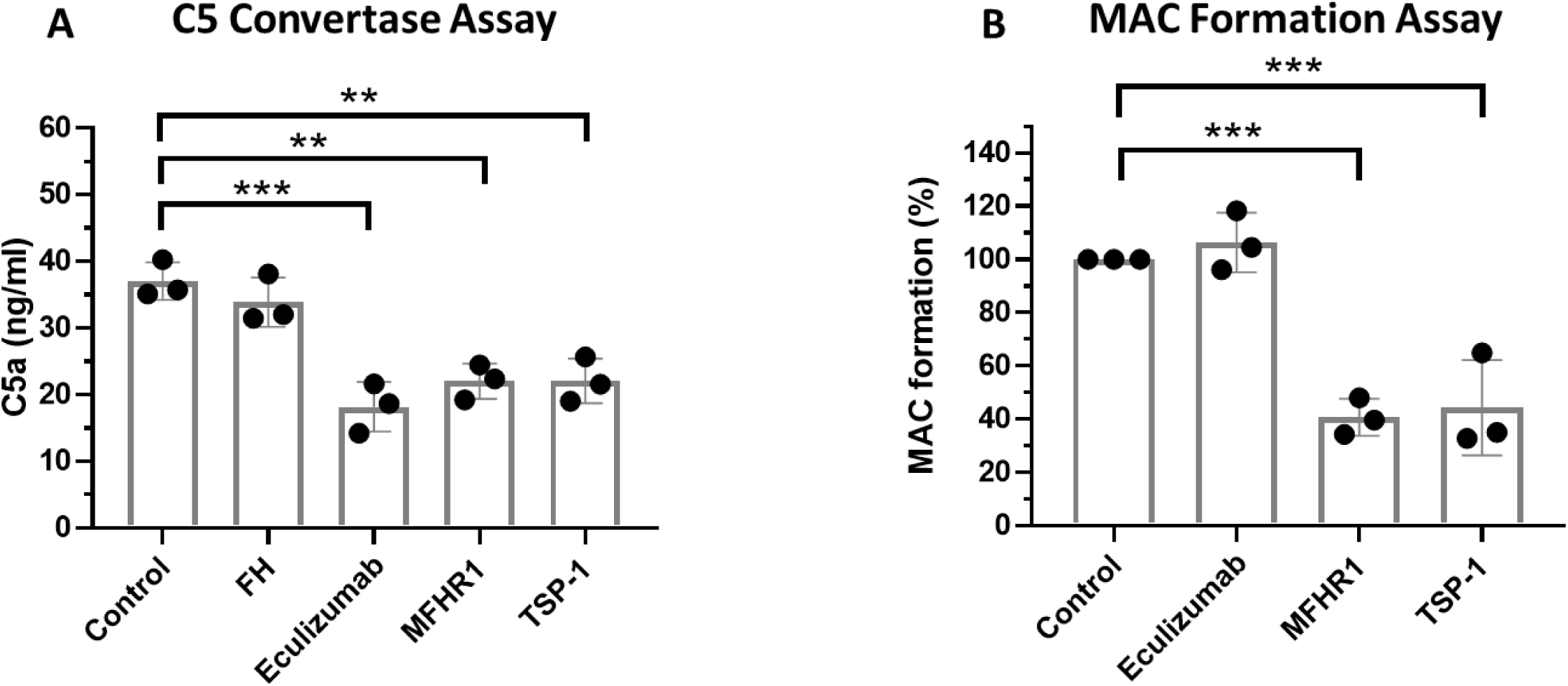
TSP-1 modulates complement at the C5 level of the complement cascade. **(A)** TSP-1 inhibits cleavage of C5. Cobra venom factor (CVF) convertase (CVFBb) was generated and added to C5 preincubated with FH, Eculizumab, MFHR1 or TSP-1. The amount of released C5a was quantified by ELISA. Results are shown as mean ± SD. *P ≤ 0.05, **P ≤ 0.01, ANOVA using Dunnett’s multiple comparison test. **(B)** TSP-1 inhibits the formation of the MAC. Sheep Erythrocytes were premixed with C7 (9 nM), C8 (7 nM) and C9 (15 nM). BSA, Eculizumab, MFHR1 or TSP-1 (1.3 µM each) were preincubated with C5b-6 (0.7 nM) and then added to the erythrocytes. Bars represent means ± SD of 3 independent experiments, **P ≤ 0.01, ***P ≤ 0.01, One-way ANOVA with Tukey’s post hoc test for comparison against control.

### TSP-1 prevents C3 deposition on PNH erythrocytes

To study the significance of the complement-regulating functions of TSP-1 demonstrated *in vitro*, we used blood samples obtained from patients with primary complement overactivation due to various primary complement-mediated diseases. The ability of TSP-1 to affect complement deposition on PNH erythrocytes was investigated in red blood cells from three PNH patients. After incubation with acid-activated AB0-adapted normal human sera (aNHS), the resulting complement-mediated damage to erythrocytes was analyzed. C3 deposition on erythrocytes was determined by flow cytometry. In acid-activated human serum, CD59-negative cells are destroyed over time by complement activation and only few cells with C3 deposits could be detected (Figure 4A). Treatment with Eculizumab inhibited erythrocyte lysis, but the cells retained C3 deposits on their surface. However, treatment of PNH erythrocytes with Eculizumab in combination with TSP-1 or Pegcetacoplan led to a significant reduction in C3-positive erythrocytes, indicating that TSP-1 is able to largely prevent complement deposition on PNH erythrocytes (Figure 4B).

**Figure 4:**
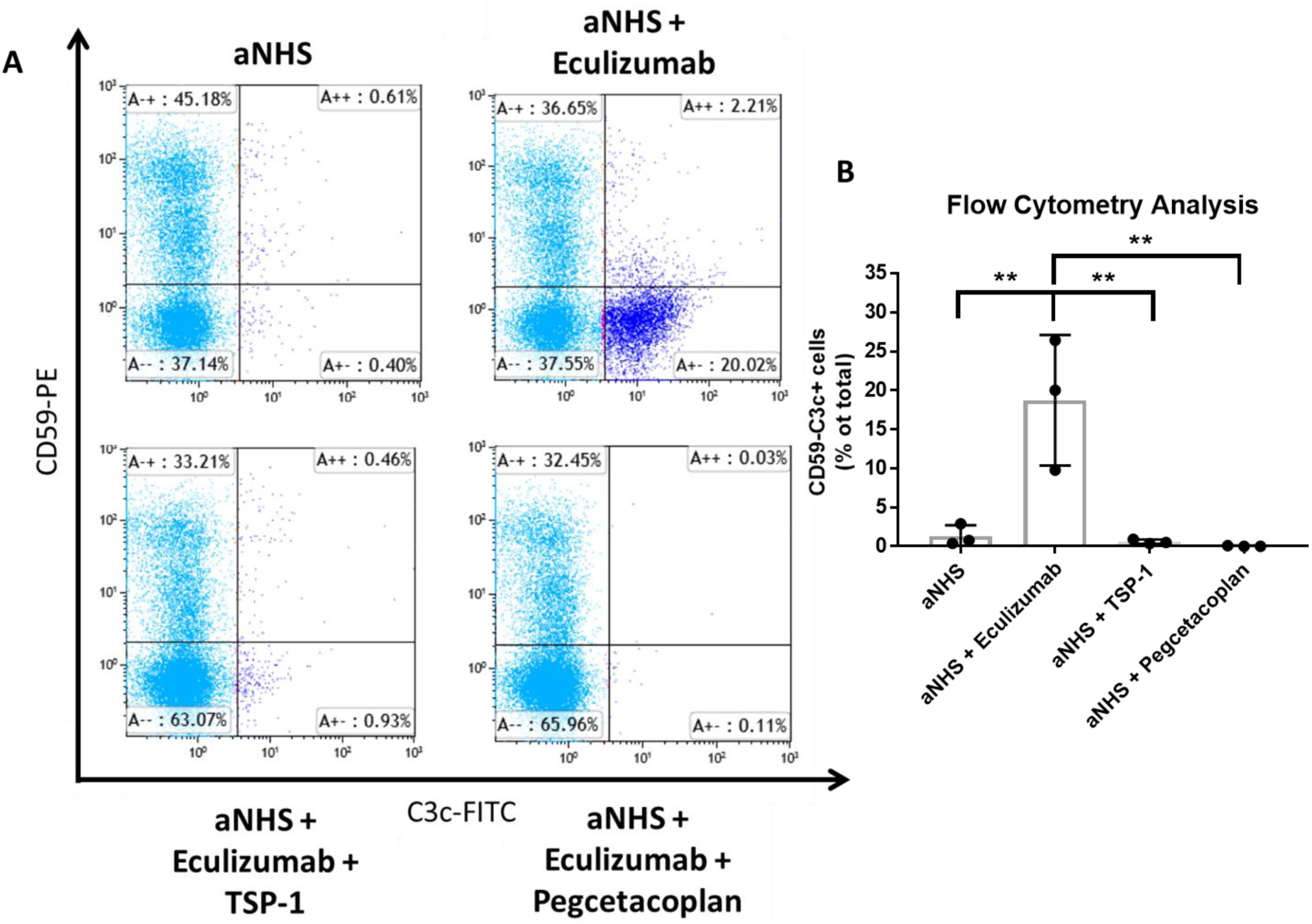
TSP-1 protects PNH erythrocytes from complement mediated opsonization. **(A)** TSP-1 prevents C3 deposition on PNH erythrocytes. PNH Erythrocytes were incubated with acidified serum alone, Eculizumab or with a combination of Eculizumab and TSP-1 or the C3 inhibitor Pegcetacoplan. Erythrocytes were stained for CD59 and C3 and percentages of positive and negative stained cells analyzed by flow cytometry. Eculizumab prevents lysis of PNH erythrocytes but leaves C3 depositions on the surface of CD59 negative erythrocytes. Combination of Eculizumab with TSP-1 or Pegcetacoplan prevents C3 deposition on CD59 negative PNH erythrocytes. **(B)** Analysis of C3 deposition on PNH erythrocytes from three different patients’ samples. Combined treatment of erythrocytes with Eculizumab and TSP-1 or Eculizumab and Pegcetacoplan significantly reduced the amount of C3 positive CD59 negative cells compared to Eculizumab treatment alone. Bars represent means ± SD of 3 independent experiments, **P ≤ 0.01, One-way ANOVA with Tukey’s post hoc test. aNHS – acidified normal human serum.

### TSP-1 inhibits hemolytic activity and C3 deposition on endothelial cells in aHUS sera

Serum from a healthy sibling (aHUS1) belonging to an aHUS family and carrying a known FH mutation for aHUS (R1215Q, SCR20) (patients details in supplemental Table S1A) was used in a hemolytic assay. The FH mutation causes strong hemolytic activity when incubated with sheep erythrocytes. TSP-1, in contrast to TSP-5, was able to reduce hemolysis when added to the aHUS1 serum, as was FH. More than 70 % of hemolysis could be inhibited at concentrations of around 200 nM TSP-1 (Figure 5A).

**Figure 5:**
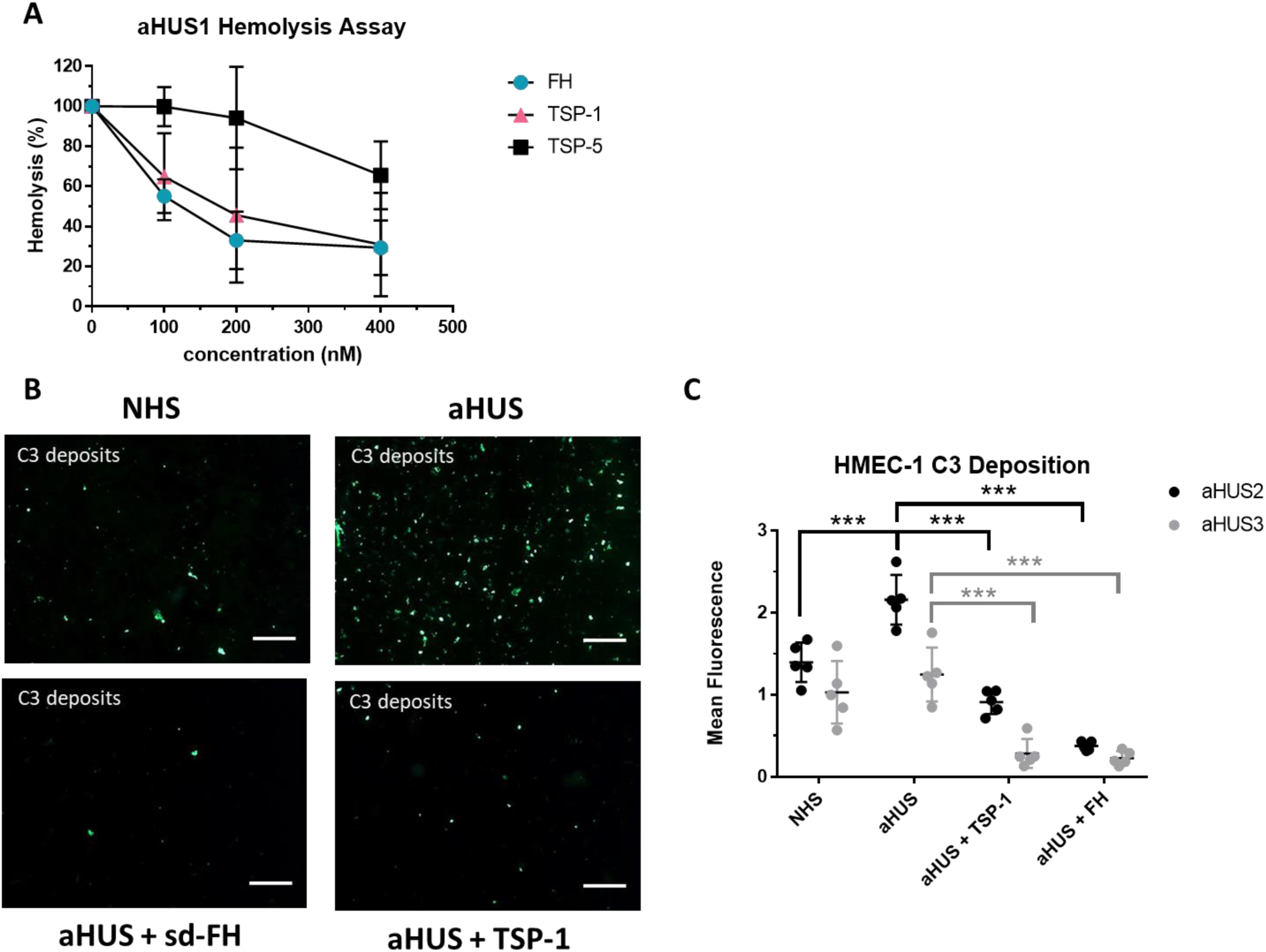
TSP-1 inhibits hemolytic activity and pathogenic C3 deposition on endothelial cells when added to the serum of aHUS patients. **(A)** TSP-1 protects sheep erythrocytes from complement induced lysis in aHUS1 serum. Sheep erythrocytes were incubated with aHUS1 serum and increasing concentrations of FH, TSP-1 or TSP-5. Data was normalized against erythrocytes treated with aHUS1 serum without inhibitors. Bars represent means ± SD of 3 independent experiments. **(B)** Representative fluorescence images of C3 deposits on HMEC-1 cells treated with aHUS sera. HMEC-1 cells were activated with ADP and incubated with 50% normal human serum or aHUS serum with or without 1µM TSP-1 or FH and stained for C3 deposits. **(C)** TSP-1 prevents C3 deposition on endothelial cells treated with aHUS sera. Mean fluorescence analysis shows that aHUS2 and aHUS 3 serum causes strong deposition of C3 molecules on HMEC-1, which could be prevented by addition of either FH or TSP-1 into the serum. C3 fluorescence intensity was measured in at least 5 randomly chosen high power fields. Results are shown as mean ± SD. ***P ≤ 0.01, One-way ANOVA using Turkey’s multiple comparison test. Scale Bar = 100 µl

Endothelial injury and complement deposition on endothelial cells are a hallmark of the pathology of complement-mediated diseases like aHUS. To investigate whether TSP-1 could influence complement deposition on endothelial cells, a cell-based assay involving human microvascular endothelial cells (HMEC-1) was performed (31). In this assay, sera from aHUS patients were incubated on ADP-stimulated HUVECs and the amount of C3 deposition was analyzed by immunofluorescence staining. The use of sera from two patients with known aHUS characteristics (aHUS2 and aHUS3, supplemental Table S1A) resulted in increased C3 deposition on HUVECs compared to normal human serum. In contrast, supplementation of the sera with TSP-1 or FH led to a significant reduction in C3 deposition (Figure 5B, 5C).

### Knockdown of TSP-1 significantly increases deposition of C3 on stimulated endothelial cells

Given that TSP-1 is stored in the Weibel-Palade bodies of endothelial cells, we investigated the effect of TSP-1 deficiency on complement deposition. HUVECs were treated with TSP-1 siRNA and stimulated with histamine as a model for endothelial injury under serum-free conditions. The siRNA treatment resulted in a reduction of approximately 40-60% in the amount of TSP-1 in HUVECs, indicating a sufficient level of suppression. Knockdown of TSP-1 significantly increased the deposition of C3 on stimulated HUVEC surfaces. However, the addition of TSP-1 could remedy the increase in C3 levels (Figure 6).

**Figure 6:**
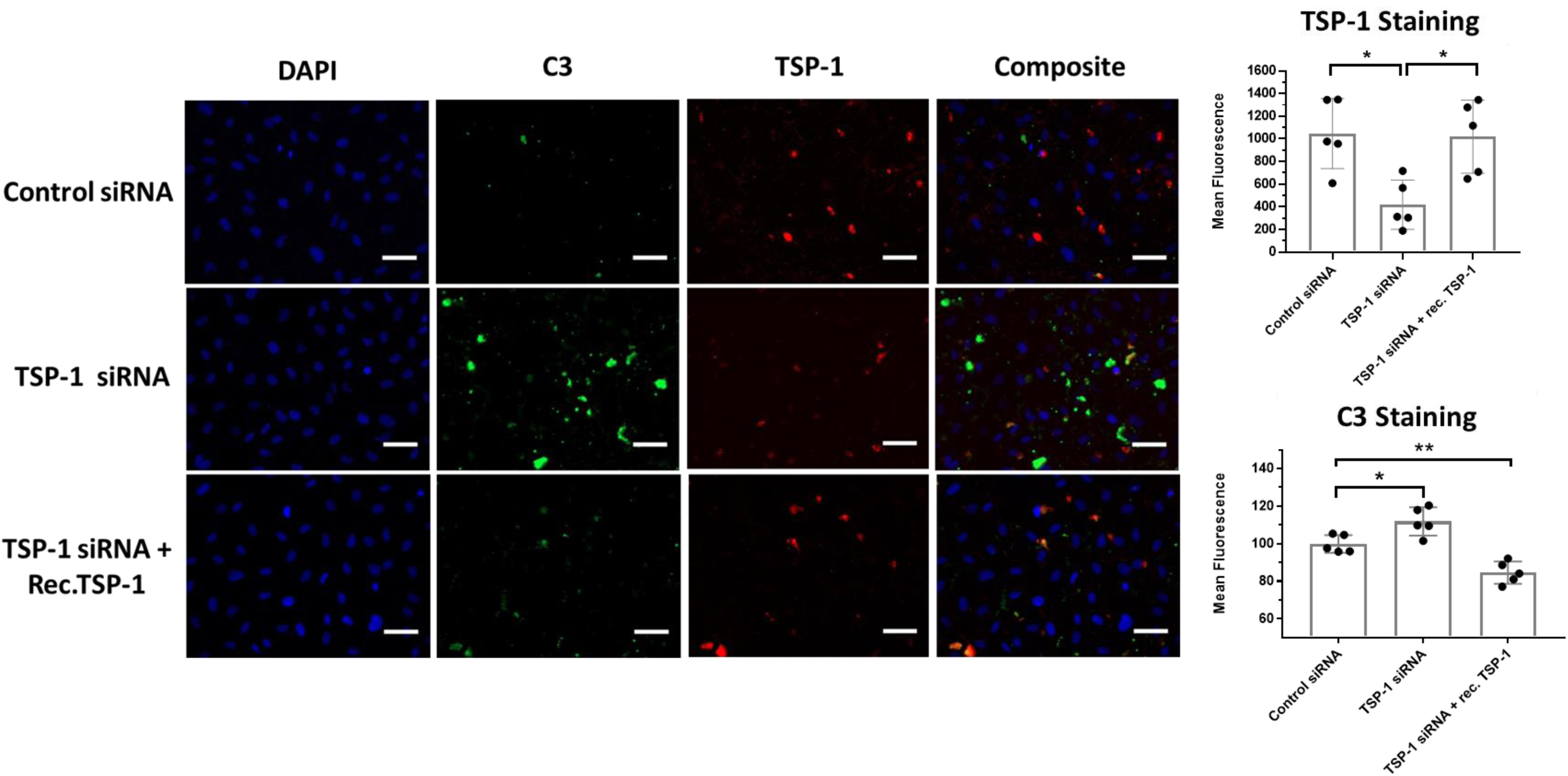
Potential role of TSP-1 in regulating complement activity on endothelial cells. Knockdown of TSP-1 increases C3 deposition on HUVECs. Left side: Representative fluorescence images of siRNA-treated HUVECs. Cells were transfected with control siRNA or TSP-1 siRNA and stimulated with histamine to mimic inflammation. In parallel, recombinant TSP-1 (rec. TSP-1) was supplemented into TSP-1 siRNA-treated cell supernatants. After 45 minutes of incubation, cells were fixed and stained for DAPI (blue), C3 (green), or TSP-1 (red). Right side: Analysis of TSP-1 and C3 staining. TSP-1 siRNA-treated samples exhibited a significant decrease in TSP-1 staining, which was reinstated by the addition of TSP-1 to the supernatant. In parallel, TSP-1 knockdown led to a substantial increase in C3 deposits on HUVECs, and this effect was ameliorated by supplementing the supernatant with TSP-1. Fluorescence intensity was measured in 5 randomly chosen high-power fields. Results are shown as mean ± SD. *P≤ 0.05, **P ≤ 0.01, One-way ANOVA using Turkey’s multiple comparison test. Scale Bar = 100 µm.

### Stimulation of HUVECs by spike protein leads to co-localization of TSP-1 with vWF, FH and C3

Stimulation of endothelial cells by the S1 spike protein has been described to promote complement activation and thrombus formation in a microfuidic assay (32). To test whether TSP-1 could play a role in complement regulation in this context, HUVECs were treated with or without 5 µg of S1 protein. Following stimulation, the cells were fixed and stained for TSP-1, C3, vWF and FH. Under normal conditions, both TSP-1 and vWF exhibited co-localization without C3 or FH on the surface of the endothelial cells (Figure 7). After stimulation, a strong deposition of C3 and FH was observed on vWF fibers in conjunction with TSP-1. Of note, the staining of TSP-1 shows higher correlation with C3 and FH staining than vWF, indicating that TSP-1 is the main mediator of this interaction.

**Figure 7:**
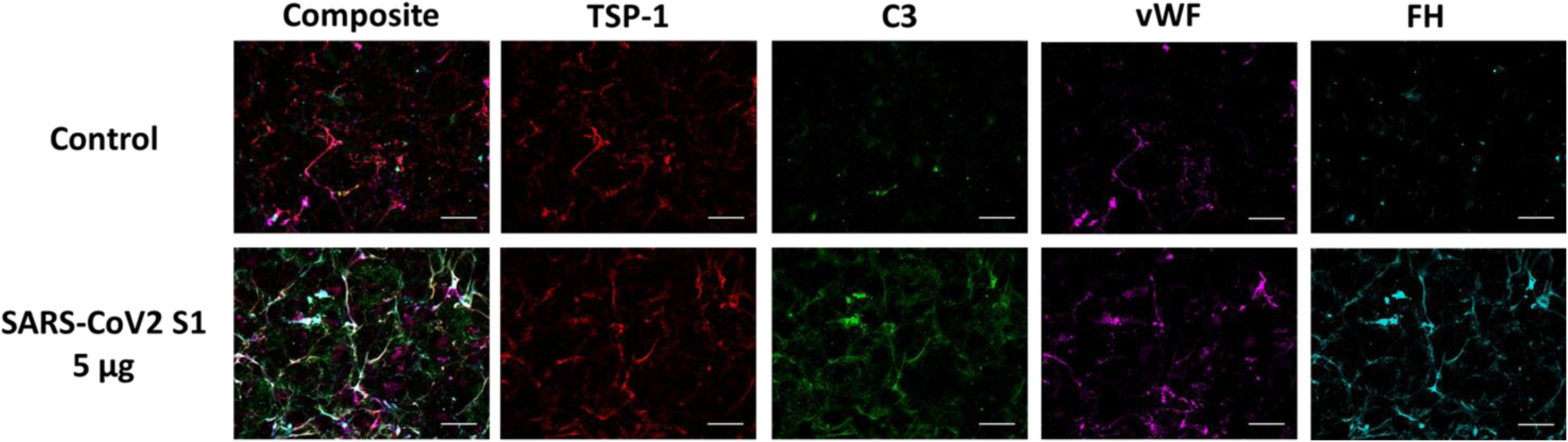
Potential role of TSP-1 in SARS-CoV-2 endothelial inflammation. SARS-CoV-2 mediated inflammation demonstrates the interplay between TSP-1, vWF and the complement system on endothelial cells. HUVECs were stimulated with hirudin treated blood, in the presence or absence of 5 µg of SARS-CoV-2 S1 protein, for 45 minutes. Following fixation, immunostaining was performed to visualize TSP-1 (red), C3 (green), vWF (magenta), and FH (cyan). In untreated samples, co-localization of TSP-1 and vWF was observed with limited involvement of C3 and FH. However, stimulation with S1 protein resulted in a significant increase in TSP-1/vWF fibers, accompanied by robust co-localized deposition of C3 and FH. Scale Bar = 50 µm.

### Significant TSP-1 staining in ANCA-vasculitis renal biopsies

Recent publications have reported significantly increased TSP-1 plasma levels in ANCA patients (33, 34). Considering that complement activation is a driving force of neutrophil extracellular trap (NET) release, we elucidated whether the complement-regulating function of TSP-1 could potentially influence this process. To evaluate a (patho-)physiological involvement of TSP-1 in ANCA-vasculitis, we stained kidney biopsies from four different ANCA-vasculitis patients (patient details in supplement Table S1C). TSP-1 showed strong positivity in all glomerular crescent lesions observed in ANCA-vasculitis biopsies (Figure 8). This was in contrast to nephrectomy controls and biopsies from patients with focal segmental sclerosis (FSGS), who also had renal impairment, hematuria and proteinuria.

**Figure 8:**
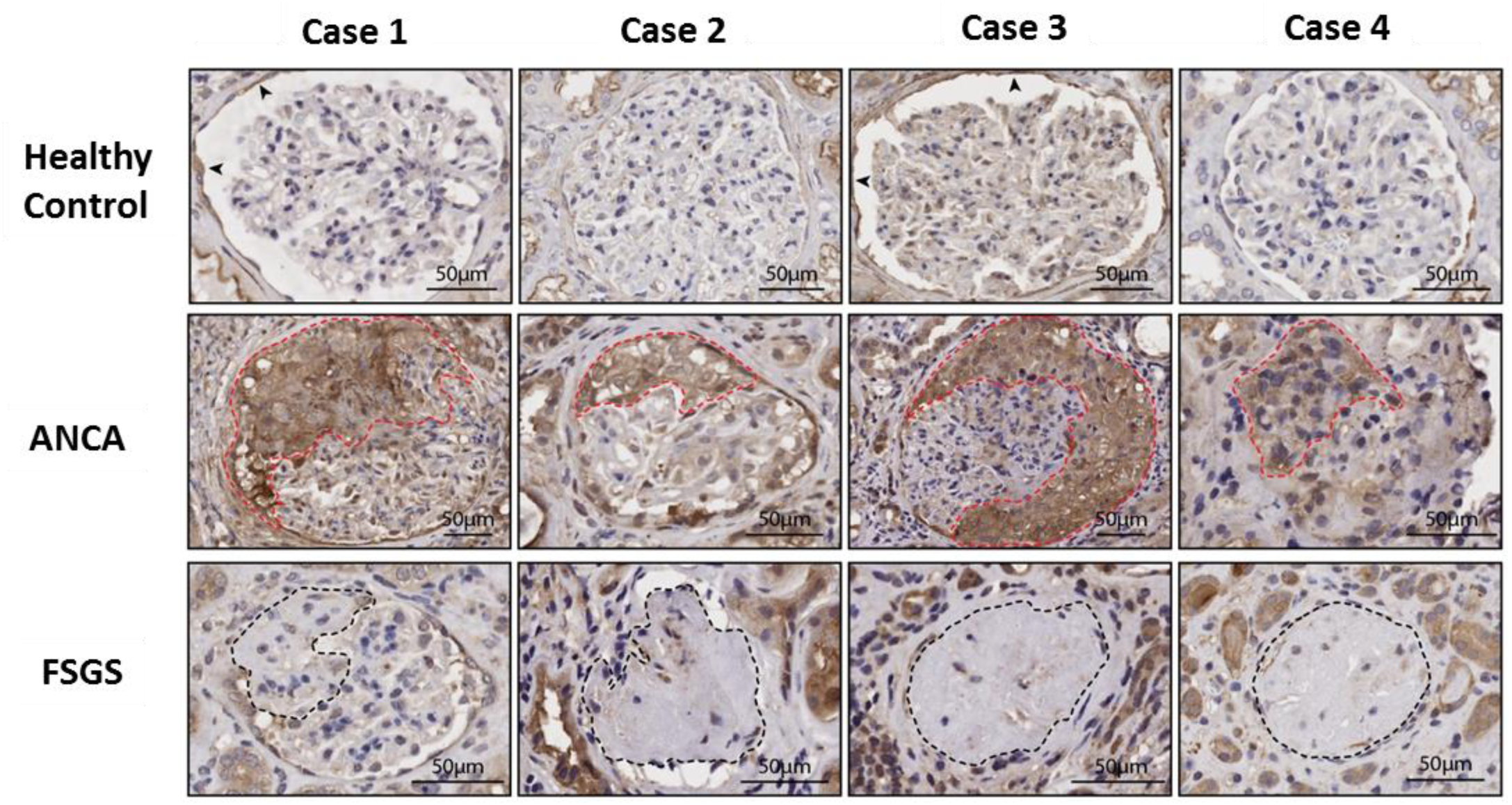
TSP-1 is involved in the pathogenesis of ANCA vasculitis. TSP-1 is significantly increased in crescents of ANCA-vasculitis patients. Representative images of TSP-1 IHC staining of kidney biopsies and cancer nephrectomies. Pronounced TSP-1 staining can be seen in glomerular crescents (red dashed lines) of four ANCA-vasculitis patients (ANCA) while TSP-1 is absent in healthy controls or patients with focal segmental glomerulosclerosis (FSGS; black dashed lines indicate sclerosis). Scale Bar = 50 µm.

### Blockade of TSP-1 in whole blood causes pronounced release of NETs in a microfluidic in vitro model of ANCA-vasculitis

To investigate the potential influence of TSP-1 on NETosis, a microfluidic experiment was conducted. PR3 antibodies isolated from an ANCA-vasculitis patient (ANCA1) were mixed with healthy hirudin-treated whole blood at a concentration of 250U/ml, comparable to the level measured in the patient. The PR3-treated blood was perfused in a microfluidic chamber coated with HUVECs. DAPI staining was used to detect DNA NETs, and NETs release was monitored through live cell imaging over a 4-hour period. Addition of ANCA-PR3 antibodies alone did not result in the visual detection of NETs (Figure 9A), as an external trigger would typically be necessary to activate neutrophil trap release (35). However, when TSP-1 was impaired by addition of TSP-1 antibody, a significant amount of NETs became detectable over the course of the experiment (see white arrows). In contrast, the addition of a mouse IgG1 isotype control antibody did not induce NET release.

**Figure 9:**
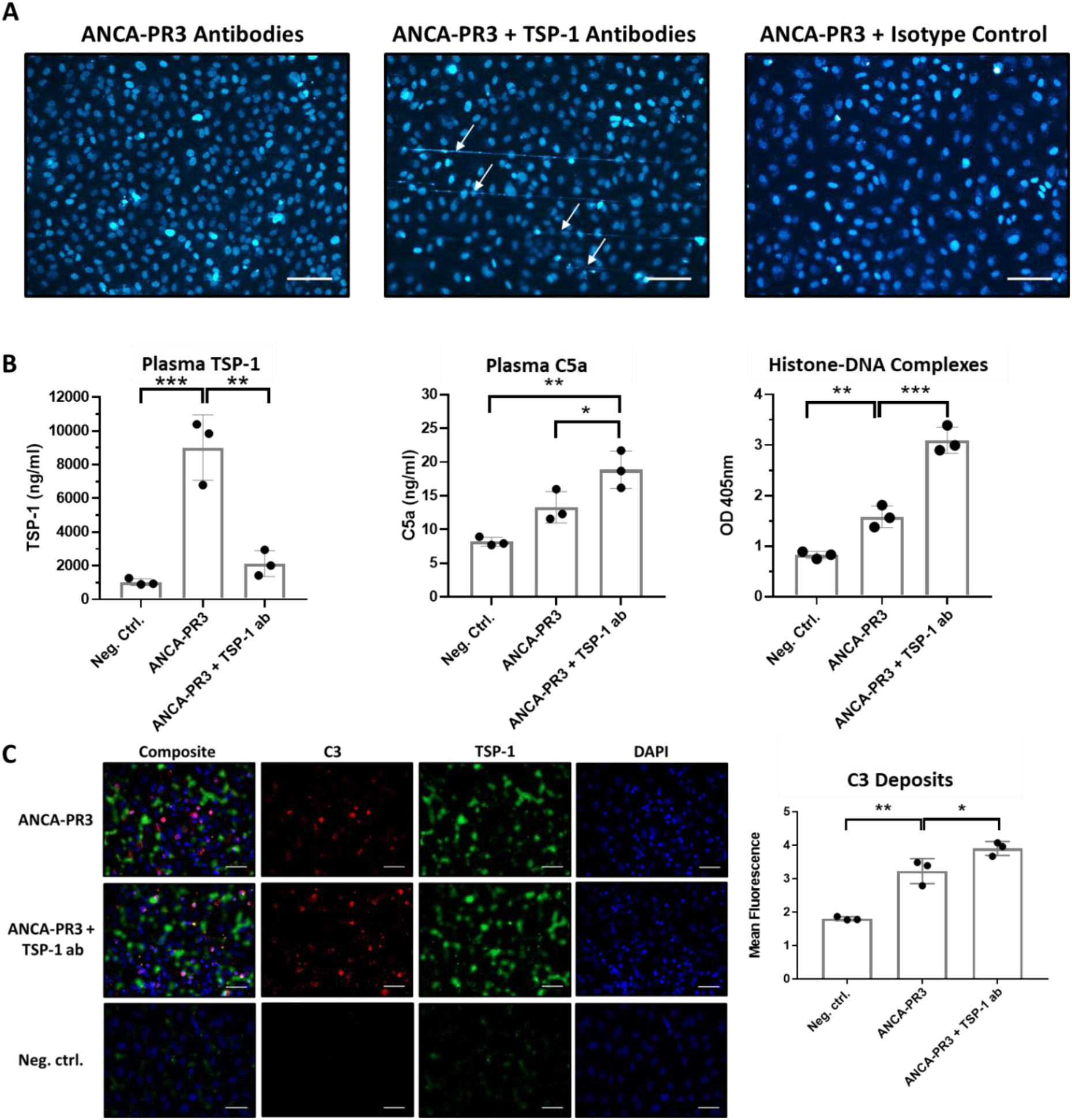
TSP-1 regulates complement activity as well as neutrophil extracellular trap release in an ANCA-vasculitis model. **(A)** Blockade of TSP-1 in ANCA-PR3 treated whole blood causes pronounced release of DNA NETs. HUVECs coated µ-slides were perfused with whole blood treated with hirudin and with ANCA-PR3 antibodies isolated from an ANCA-vasculitis patient (patient ANCA1). Whole blood was stained with DAPI and NET release monitored for 4 hours via live cell imaging. Treatment of whole blood did not cause visible NET release. ANCA-PR3 antibodies in combination with TSP-1 blockade caused pronounced NET release on endothelial cells. In contrast, ANCA-PR3 antibodies in combination with a TSP-1 isotype control antibody did not have an effect. Scale Bar = 100 µm **(B)** Antibody mediated blockade of TSP-1 modulates plasma C5a levels as well as the release of histone-DNA complexes in a microfluidic experiment mimicking ANCA-vasculitis. ANCA-PR3 antibodies induce a significant increase of plasma TSP-1 levels which can be suppressed by addition of a TSP-1 antibody. The addition of ANCA-PR3 antibodies to whole blood leads to a substantial increase in plasma C5a levels compared to untreated whole blood (Neg. ctrl.). This increase is further amplified when combined with a TSP-1 antibody. Treatment of whole blood with ANCA-PR3 antibodies results in a significant rise in released histone-DNA complexes. This increase is further intensified when TSP-1 antibody is added. Bars represent means ± SD of 3 independent experiments. *P ≤ 0.05, **P ≤ 0.01, ***P ≤ 0,001, One-way ANOVA using Turkey’s multiple comparison test. **(C)** Blockade of TSP-1 causes increased C3 deposition on HUVECs perfused with ANCA treated whole blood. Treatment of whole blood with ANCA-PR3 causes a notable enhancement of both C3 and TSP-1 deposition on HUVECs compared to untreated samples (Neg. ctrl.). Furthermore, the increase in C3 deposition is significantly augmented when combined with a TSP-1 antibody. Mean fluorescence analysis of C3 staining is presented on the right. Bars represent means ± SD of 3 independent experiments. *P ≤ 0.05, ***P ≤ 0.01, One-way ANOVA using Turkey’s multiple comparison test. Scale Bar = 50 µm.

To further elucidate these findings, the experiments were repeated including the collection of revealed a 9-fold increase in TSP-1 levels, a 1.9-fold increase in C5a and a 1.75-fold increase in histone-DNA complexes compared to negative control (Figure 9B). In samples treated with TSP-1 antibody, TSP-1 levels were only increased 2-fold, while C5a levels increased by 2.5-fold and histone-DNA complexes by 3.7-fold compared to negative control. On HUVECs, ANCA-PR3 antibodies led to a pronounced increase in TSP-1 staining, correlating with plasma TSP-1 levels (Figure 9C). In addition, antibody-mediated blockade of TSP-1 resulted in a significant increase in C3 deposition on HUVECs.

## Discussion

Taken together, our results revealed that TSP-1 acts as an inhibitor of alternative complement pathway activation. We uncovered new interaction partners of TSP-1, providing insights into the mechanisms of complement regulation by this versatile protein. TSP-1 shows the ability to impede AP activation at both the C3 and C5 levels. This inhibitory effect extends to the prevention of C3 and C5 cleavage and effectively hinders the formation of the MAC (summarized in Figure 10).

**Figure 10:**
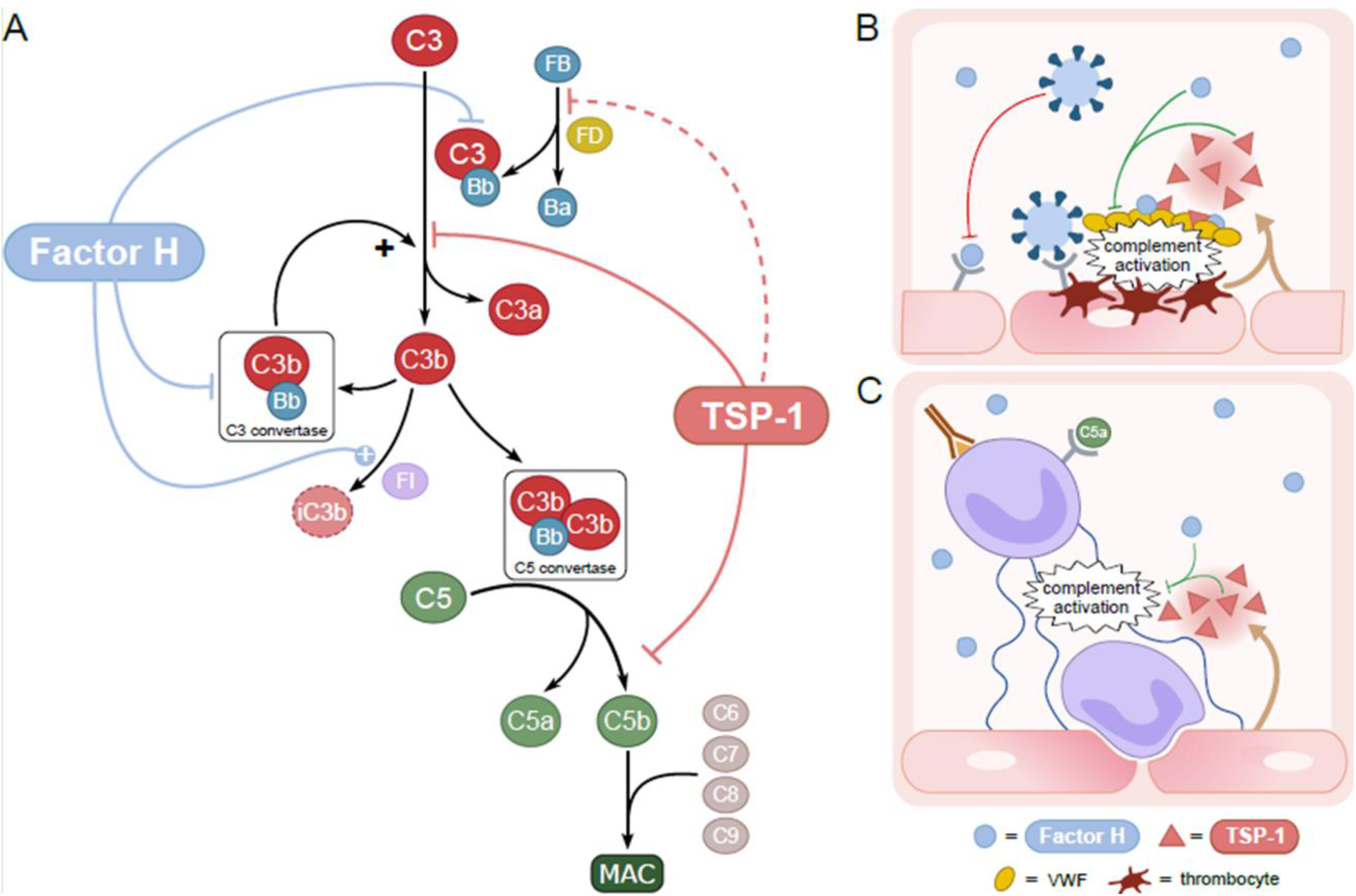
Summary of TSP-1 effects on the alternative pathway and physiological significance. **(A)** Complement inhibitory functions of TSP-1 in comparison to factor H are schematically illustrated: TSP-1 prevents cleavage of FB by FD in vitro, although binding to FB has not been confirmed by SPR and therefore not physiologically relevant (dashed line). TSP-1 binds to C3 and prevents the cleavage of C3 into C3a and C3b. Furthermore, TSP-1 binds to C5 and prevents its cleavage into C5a and C5b and the formation of the MAC. Possible physiological and pathophysiological complement regulatory functions of TSP-1 in simplified models: secondary complement activation occurs in SARS-CoV-2 hyperinflammation and thrombosis **(B)** due to COVID spike protein induced endothelial impairment and thrombotic events related to the formation of ultra large vWF multimers and in ANCA-vasculitis **(C)** due to neutrophil activation, netosis and subsequent vasculitis. In these local overwhelming conditions, inhibition by FH might not be sufficient to control complement activation. Therefore, additional complement inhibitory functions by locally released TSP-1 from endothelia and/or thrombocytes could be physiologically relevant regarding control of excessive complement activation especially on surfaces.

To confirm TSP-1-dependent complement regulation in a more physiological context, we used various sera and erythrocytes from patients with primary complement-mediated diseases in *ex vivo* assays. In aHUS sera, TSP-1 was able to prevent complement activation and subsequent hemolysis of sheep erythrocytes and inhibited the deposition of C3 on the surface of endothelial cells. On PNH erythrocytes, the addition of TSP-1 prevented C3 deposition. These experiments demonstrate that TSP-1 can influence and prevent complement activation under different pathological conditions.

The complement regulatory role of TSP-1 certainly differs from that of FH. Unlike FH knockout mice, which develop primary complement-related disorders resembling C3 glomerulopathy, the knockout of TSP-1 in mice does not lead to such disorders. However, the lack of TSP-1 mutations in humans is explained due to loss-intolerance during human development (36, 37). FH is abundantly present in the bloodstream and is able to control complement activation at initiation, especially on host surfaces (38). In contrast, TSP-1 is only found in small amounts in the fluid phase but can be rapidly released from activated endothelial cells and/or platelets (2, 3). In inflammatory conditions, TSP-1 might act locally as a complement regulator, either alongside or in combination with FH. It is noteworthy that the complement-regulating properties of TSP-1 are functionally distinct from those of FH (Figure 7), rather implying a collaborative role for these two complement inhibitors. Moreover, TSP-1 does not influence decay acceleration activity of FH and has no intrinsic cofactor activity (supplemental Figure 2A, 2B).

The local complement inhibitory effect of TSP-1 may have physiological relevance in endothelial stimulation scenarios, such as SARS-CoV-2-related hyperinflammation/thrombosis or complement-associated vasculitis diseases like ANCA-vasculitis (21, 22). In SARS-CoV-2-related hyperinflammation/thrombosis, complement involvement is thought to be triggered by two mechanisms: endothelial damage by the spike protein and the appearance of large vWF multimers associated with complement activation and a severe disease process (23, 24). Besides complement activation elevated TSP-1 levels are found in severe Covid-19 as well as long Covid patients (39, 40).

TSP-1 is present in Weibel Palade bodies of endothelial cells in conjunction with vWF. Both proteins are released from endothelial cells upon stimulation as in vasculitis (7). TSP-1 co-localizes with vWF and previous studies have also demonstrated the recruitment of complement proteins from the AP to vWF molecules (41, 42). While vWF has been implicated in complement-mediated diseases and thrombosis, its exact role in relation to complement still remains unclear (20). Licht et al. (43) described increased damage to blood outgrowth endothelial cells lacking vWF, but Norris et al. (44) instead showed that the absence of vWF protects human microvascular endothelial cells from complement-mediated damage (43, 44). In our experiments, the knockdown of TSP-1 in CD31-stimulated endothelial cells led to an increased amount of C3 on the surface of endothelial cells, which could be reversed by addition of recombinant TSP-1. Stimulating HUVECs with SARS-CoV-2 spike protein induced thrombosis (32). We could demonstrated a clear co-localization of TSP-1, FH and C3 on vWF fibers. Hence, the local complement inhibitory role of TSP-1 could play a crucial role in containing complement activation, particularly in situations where broad inhibition, such as that provided by FH, proves inadequate in managing highly localized and intensified processes (as depicted in Figure 7B). It is also conceivable that TSP-1 aids FH by recruiting it to cell surfaces or vWF.

In ANCA-vasculitis, complement activation and NETosis are prominent drivers of vascular inflammation and injury (27). Complement component C5a is capable of triggering NETosis, whereupon the formed NETs serve as a platform for complement activation. Under these circumstances, locally released TSP-1 may additionally help to prevent excessive complement activation in this detrimental cycle (Figure 7C). Consistent with this, significantly elevated plasma TSP-1 levels have recently been demonstrated in ANCA-vasculitis patients (657.47 ± 64.17 pg/ml) compared to healthy controls (253.78 ± 37.10 pg/ml, P ≤ 0.005) (33, 34).

In accordance with this, our current findings detected prominent TSP-1 staining in renal biopsies of ANCA-vasculitis patients in glomerular crescents. Additionally, depletion of TSP-1 in an ANCA-vasculitis model using patients’ PR3 antibodies led to a distinct increase in complement activation and NETosis, supporting the influence of TSP-1 on complement and disease activity in ANCA-vasculitis. In this study, we demonstrate the complement inhibitory function of TSP-1 in human blood samples from healthy individuals and from patients with complement-related diseases, as well as in serum free conditions performing various assays. We initially used heparin-free TSP-1 with an N-terminal histidine tag, which could be a source of error, since TSP-1 can bind to histidine-rich proteins, potentially causing unspecific aggregation (45). To address this issue, we used untagged TSP-1 purified from human platelets and repeated our main findings (Figure 1B, supplemental Figure S1). The complement regulatory functions of platelet-isolated TSP-1 remained consistent in our assays, excluding unspecific protein aggregation as the cause of complement regulation.

While TSP-1 inhibits FD mediated cleavage of FB in vitro, SPR measurement detected only weak binding to FB, suggesting that FB cleavage might not be essential under physiological conditions. In contrast, strong binding in SPR was confirmed for C3, C5, C8 and FH, suggesting that the TSP-1 effects on C3 and C5 cleavage and MAC formation are physiologically relevant processes.

TSP-1 is known to have various physiological effects stemming from interactions with numerous receptors and other proteins (37, 46). Therefore, in the more complex *in vivo* setting, complement regulation by TSP-1 may be additionally influenced by other effector pathways of TSP-1. However, by employing different model systems of various complement-mediated diseases, we were able to show a consistent inhibitory effect of TSP-1 on the alternative complement pathway.

Overall, our findings provide strong evidence that TSP-1 acts as a novel complement inhibitor. While complement regulation by TSP-1 appears less relevant under physiological conditions, it seems to be an important co-player in the context of pronounced endothelial inflammation. Hence, it represents a new mechanism in the intricate interplay between endothelia, complement and platelets. The regulatory effect especially gains significance in secondary complement-mediated vasculitis disease, where additional complement-inhibitory function is required locally due to excessive complement activating processes (Figure 10B/C). Given its inhibitory effect, particularly on surface complement activation, and its interaction with vWF, it is intriguing to hypothesize that TSP-1 protects endothelial cells from complement mediated damage.

Considering that complement activation itself can contribute to vasculitis or thrombotic conditions, TSP-1 may also be involved in mitigating the sequelae of an overactive complement system, further preventing anaphylatoxin C3a and C5a formation (18, 19, 47). Therefore, our findings may contribute to an improved understanding of these complex pathogenic mechanisms and might pave the way for the development of novel therapeutic strategies.

## Methods

### Sex as biological variable

Sex was not considered as a biological variable. However, blood and tissue samples examined in our study were from both sexes. Patient details are summarized in the supplement.

### Generation and purification of Thrombospondin-1

Human TSP-1 cDNA was obtained from Sinobiological (HG10508-UT) and subcloned into a pFastbac1 vector containing a gp67 signal peptide for secretion and an N-terminal 10xHis tag for purification. Bacmids were generated using the Bac-to-Bac vector kit (ThermoFisher, 10360014) and used to generate recombinant baculovirus. TSP-1 was expressed in Spodoptera frugiperda 9 (ThermoFisher, 11496015) cells and purified from cell supernatants via Ni-NTA columns. TSP-1 concentrations were determined by ELISA (R&D, DY3074).

### Proteins and Sera

Complement proteins (FB, FD, FH, FI, C3, C3b, C5, C5b6, C6, C7, C8, and C9) as well as FH-deficient serum were obtained from Complement Technologies. Cobra venom factor (CVF) was purchased from Quidel, and Eculizumab (Soliris) as well as Pegcetacoplan (Aspaveli) was obtained from remnants of infusions. TSP-5 was obtained from R&D. Serum samples from healthy donors or patients with aHUS as well as EDTA treated whole blood from PNH patients were obtained following standard procedures (48). PR3 antibodies from ANCA1 patient were obtained by purifying plasma using a protein G affinity column (Cytiva) following manufacturer’s instructions. PR3 concentrations were determined with an Algeria ELISA analyzer.

### Alternative Pathway ELISA

Alternative pathway activity was measured using commercially available ELISA kits following manufacturer’s instructions (Avar Life Science, COMPLAP330). C5a was quantified by C5a ELISA (Abcam, ab193695) according to the manufactures protocol with the following modification: during incubation period 30 µl of ice cold 20 mM EDTA was added to the samples. Cell death detection ELISA (Roche, 11544675001) was used to detect histones H1, H2a, H2b, H3 and H4, single and double stranded DNA from microfluidic experiments according to manufacturer’s instructions.

### Hemolytic Assay

Sheep erythrocytes (sE) (Fiebig) were washed and diluted with GVB buffer containing MgEGTA and the experiment performed as described before (49). Released hemoglobin from erythrocytes was measured via absorption measurement at 414 nm.

### TSP-1 binding ELISA to complement proteins

Complement proteins FH, FB, C3, C5, C6, C7, C8, C9 or BSA were coated on microtiter plates at a concentration of 133 nM each in PBS overnight at 4 °C on Nunc maxisorb 96 well plates. Unbound proteins were washed with PBS containing 0.05 % Tween20 and the wells blocked with PBS containing 2 % BSA for 1 h at room temperature. After washing, recombinant TSP-1 was added to the wells at 10 µg/ml and incubated for 2 h at room temperature. Bound TSP-1 was detected using a mouse anti-human TSP-1 (Merck, MABT879) antibody for 2 h at room temperature, followed by a sheep anti-mouse HRP (GE Healthcare, NA931V) secondary antibody for 1 h at room temperature. Colorimetric detection was performed using TMB substrate. The reaction was stopped after 10 min and optical density at 450 nm measured

### SPR Biacore Measurements

Kinetic analysis of the interaction between TSP-1 and complement proteins was performed on a BIAcore 3000 instrument (Biacore AB, Uppsala, Sweden). TSP-1 was bound to CM5 chips at a concentration of 0.1 µM via covalent bond. Binding to complement proteins FH, FB, C3, C3b, C5 and C8 was assessed over a range of concentrations (12.3, 37.03, 111.1, 333.3, 1000 nM) in PBS + 0.05% Tween20. Binding data were fitted to a 1:1 Langmuir binding model and on and off rates determined for calculating affinity constants. Between experiments chip surface was regenerated with 10 mM sodium hydroxide.

### FB cleavage Assay

The ability of TSP-1 to inhibit FD mediated cleavage of FB was performed as described before (50). FB cleavage products were visualized with coomassie staining after SDS-PAGE. Uncleaved and cleaved FB band intensities were determined using ImageJ.

### C3 Convertase Assay

C3 convertase activity was assessed as described before (50). Briefly, C3(H_2_O) was generated by incubating C3 with 200 mM methylamine for 30 min at 37 °C. The convertase was generated by incubating C3(H_2_O) with FB and FD. After stopping the reaction, increasing amounts of TSP-1 and C3 were added to the mixture and the amount of generated C3α‘ visualized with a coomassie stain after SDS-PAGE. C3α‘ band intensity was determined using ImageJ.

### C5 Convertase Assay

25 nM CVF, 25 nM FB, and, 2.5 nM FD were diluted in PBS containing 5 mM MgCl_2_. The reaction was incubated for 1 hour at 37°C to build up C5 convertase (CVFBb). 200 nM of C5 was mixed with a 16-fold molar excess of test proteins and incubated for 30 minutes at room temperature. After incubation the two samples were mixed and the amount of released C5a was quantified by ELISA (Abcam).

### MAC formation assay

The ability of TSP-1 to inhibit the formation of the membrane attack complex (MAC) on the surface of sheep erythrocytes was performed as described before (49). The amount of released hemoglobin was determined by OD measurement of supernatants at 414 nm.

### Inhibition of PNH C3 deposition

The ability of TSP-1 to prevent C3 deposition on the surface of PNH erythrocytes was performed as described before (51). Briefly, EDTA blood from three different PNH patients (patient details in supplement Table S1B) was obtained and washed three times in saline. 2 µl of erythrocytes were mixed with PBS alone or 0.5 µM Eculizumab with or without 1.5 µM TSP-1 or 12 µM Pegcetacoplan in a total volume of 10 µl. 30 µl of AB0 matched, acid activated (0.1 M HCl, 1:20 diluted) normal human serum (aNHS) supplemented with 2mM MgCl_2_ was added to the cells and incubated for 24 h at 37 °C. The cells were washed with PBS containing 2 mM EDTA and stained with antibodies against CD59 (1:100, anti-human CD59 PE, Biolegend) and C3c (1:25, anti-human C3c FITC, Dako F020102) before flow cytometry analysis.

### Inhibition of C3 deposition on HMEC-1

The ability of TSP-1 to inhibit C3 deposition on endothelial cells was assessed with an assay described before (31). Briefly, HMEC-1 cells were activated with 10 µM ADP in HBSS for 10 min. Next, cells were incubated with either normal human serum or aHUS patient serum treated with or without 1 µM of FH or TSP-1 for 4 h at 37 °C, respectively. After incubation the cells were fixed in 4 % PFA and stained with rabbit anti human C3c FITC (1: 100 Dako F020102) antibody, capable of recognizing C3 and C3b, and DAPI. Mean fluorescence of obtained images was measured using ImageJ.

### siRNA mediated knockdown of TSP-1

Small interfering RNA (siRNA) to knockdown TSP-1 expression in HUVECs was obtained from Qiagen (GS7057). AllStars siRNA was used as negative control. Transfection was performed using Oligofectamine 2000 (Invitrogen) reagent in Gibco 199 serum free medium following the manufacturer’s instructions. The cells were incubated for 48 hours at 37°C before use in experiments.

On the day of experiment, HUVECs were treated with HEPES buffer containing 50µM histamine with or without TSP-1. The cells were incubated at 37 °C for 45 min, washed with HEPES and fixed with 4% PFA. The cells were blocked with 1% BSA before staining. HUVECs were stained with goat anti human C3 (1:100, Complement Technologies A213) and mouse anti human TSP-1 (1:100, Santa Cruz sc-59887) followed by donkey anti mouse Alexa 488 (1:500) and donkey α goat IgG Alexa 555 (1:500, Thermo A21432). Nuclei were stained with DAPI before mounting with mowiol. Mean fluorescence of obtained images was measured using ImageJ.

### Stimulating HUVECs with SARS-CoV19 spike protein

HUVECs were treated with hirudinated whole blood (1:1 mixed with HEPES) with or without 5 µg of SARS-CoV2 S1 protein (Biotrend, 230-01101). The cells were incubated at 37 °C for 45 min, washed with HEPES and fixed with 4% PFA. The cells were blocked with 1% BSA before staining. HUVECs were stained with goat anti human C3 (A213), rabbit anti human TSP-1 (sc-59887), rabbit anti human vWF (1:200, Dako GA527) and goat anti human FH (1:500, Complement Technologies A237). Nuclei were stained with DAPI. Mean fluorescence of obtained images was measured using ImageJ.

### Staining TSP-1 in renal biopsies

Immunohistochemistry (IHC) staining of FFPE kidney biopsy and cancer nephrectomy samples was performed applying standard procedures as previously described (52). After heat induced antigen retrieval (Citrate buffer, pH6) of samples an anit-TSP1 antibody (1:400, Merck/Millipore MABT879) was diluted in blocking solution and incubated at 4°C overnight. Samples were washed in PBS and HRP linked secondary antibody (1:500, Agilent/Dako, P0447) was diluted and applied for 30 min. Finally, samples were washed in PBS. DAB chromogen (Agilent/Dako, K3468) staining reagent was applied following the manufacturer’s instructions and counterstained using Hematoxylin. Samples were digitalized using a slide scanner (Ventana DP 200, Roche Diagnostics Deutschland GmbH).

### In vitro ANCA-vasculitis model

A microfluidic system with an air pressure based pump system (IBIDI GmbH, Munich, Germany) was used. HUVECs were first cultured on gelatin coated 0.2 Luer μ-slides (IBIDI GmbH, Munich, Germany) for 48 h prior to the experiment. For microfluidic experiments, approximately 350.000 cells per slide were used.

On the day of the experiment HUVECs were once washed with HEPES and consecutively perfused (pressure: 16.2m.bar, shear rate: 5 dyn/cm², flow rate: 0.5ml/min) with hirudinated whole blood (1:1 mixed with HEPES). Prior perfusion, hirudinated blood was treated with ANCA-PR3 antibody (250 U/ml final concentration) with or without TSP-1 antibody (A6.1, 100µg/ml final concentration) or an isotype control antibody. Additionally, DAPI was added to whole blood as a marker for live imaging of DNA NET release. Untreated hirudinated whole blood was used as negative control. After 4 hours of incubation, supernatants were centrifuged and collected for analysis of TSP-1, C5a as well as released histones and DNA. HUVECs were fixed and stained for C3 (1:200, Complement Technologies A213) and TSP-1 (1:100, Santa Cruz sc-59887) for immunofluorescence imaging.

### Statistics

All graphs and statistics were created using Prism8 (GraphPad Software). One-way ANOVA with post hoc Turkey’s multiple comparisons test was used for statistical evaluation as indicated. Statistical significance was defined as ***P ≤ 0.001, **P ≤ 0.01, and *P ≤ 0.05.

### Study approval

This study is registered at the German clinical trials register (DRKS00025182) and was approved by the ethics committee of Freiburg (No. 21-1324 and 21-1288). All patients included in this study provided written consent. Patients’ details are provided in supplemental methods (Table S1A-C).

## Data Availability

Values for all data points in graphs are reported in the Supporting Data Values file. For original data, please e-mail the corresponding author.

## Acknowledgments

We thank the Lighthouse Core Facility in Freiburg for access to their flow cytometers. The Lighthouse Core Facility is funded in part by the Medical Faculty, University of Freiburg (Project Numbers 2021/A2-Fol; 2021/B3-Fol) and the DFG (Project Number 450392965).

K.H., S.K. and J.K. are supported by Deutsche Forschungsgemeinschaft DFG grant (HA 9131/2-1). C.S. was supported by the German Research Foundation (DFG—Deutsche Forschungsgemeinschaft) (project-IDs 241702976, 438496892, 501370692), CRU329 (project-ID 386793560), SFB1453 (project-ID 431984000), SFB1160 (project-ID 256073931)) and the Wilhelm Sander-Stiftung (Wilhelm Sander Foundation).

## Author contributions

S.K., B.Z., T.T. and K.H. designed the experiments. S.K., J.K., X.L., and T.T. performed the research and collected the data. S.K., H.W. and R.G. performed SPR-analysis, M.R. and C.S. performed kidney biopsy staining for TSP-1. S.V., X.L. and C.G. performed microfluidic experiments. S.K., S.S., C.G., T.T. and K.H. analyzed the data. M.P., J.P. and K.H. provided the clinical data. S.K., S.S., E.L.D., M.P., B.Z., J.P., T.T. and K.H. wrote the manuscript which was revised by all other authors.

## Materials & Correspondence

Karsten Häffner, Innere Medizin IV, Mathildenstr. 1, 79102 Freiburg, Germany

## Key Points

TSP-1 inhibits complement activation at the C3 and C5 level, acting as a potent additive local complement regulator.

Complement regulation by locally released TSP-1 plays a relevant, additive role in vasculitis

## References

1. Adams JC, Lawler J. The thrombospondins. Cold Spring Harb Perspect Biol. 2011;3(10):a009712.

2. Dubernard V, et al. Evidence for an alpha-granular pool of the cytoskeletal protein alpha-actinin in human platelets that redistributes with the adhesive glycoprotein thrombospondin-1 during the exocytotic process. Arterioscler Thromb Vasc Biol. 1997;17(10):2293–2305.

3. Tuszynski GP, Nicosia RF. The role of thrombospondin-1 in tumor progression and angiogenesis. Bioessays. 1996;18(1):71–76.

4. Wight TN, et al. Light microscopic immunolocation of thrombospondin in human tissues. J Histochem Cytochem. 1985;33(4):295–302.

5. Leung LL. Role of thrombospondin in platelet aggregation. J Clin Invest. 1984;74(5):1764– 1772.

6. Wang A, et al. Thrombospondin-1 and ADAMTS13 competitively bind to VWF A2 and A3 domains in vitro. Thromb Res. 2010;126(4):e260–265.

7. Bonnefoy A, Hoylaerts MF. Thrombospondin-1 in von Willebrand factor function. Curr Drug Targets. 2008;9(10):822–832.

8. Murphy-Ullrich JE, Suto MJ. Thrombospondin-1 regulation of latent TGF-β activation: A therapeutic target for fibrotic disease. Matrix Biol. 2018;68–69:28–43.

9. Kale A, Rogers NM, Ghimire K. Thrombospondin-1 CD47 Signalling: From Mechanisms to Medicine. Int J Mol Sci. 2021;22(8):4062.

10. Happonen KE, et al. Regulation of complement by cartilage oligomeric matrix protein allows for a novel molecular diagnostic principle in rheumatoid arthritis. Arthritis Rheum. 2010;62(12):3574–3583.

11. Carron JA, et al. Factor H co-purifies with thrombospondin isolated from platelet secretate. Biochim Biophys Acta. 1996;1289(3):305–311.

12. Vaziri-Sani F, et al. Factor H binds to washed human platelets. J Thromb Haemost. 2005;3(1):154–162.

13. Ricklin D, Lambris JD. Complement in immune and inflammatory disorders: pathophysiological mechanisms. J Immunol. 2013;190(8):3831–3838.

14. Noris M, Remuzzi G. Atypical hemolytic-uremic syndrome. N Engl J Med. 2009;361(17):1676–1687.

15. Bessler M, et al. Paroxysmal nocturnal haemoglobinuria (PNH) is caused by somatic mutations in the PIG-A gene. EMBO J. 1994;13(1):110–117.

16. Malato A, et al. Thrombotic complications in paroxysmal nocturnal haemoglobinuria: a literature review. Blood Transfus. 2012;10(4):428–435.

17. Dzik S. Complement and Coagulation: Cross Talk Through Time. Transfus Med Rev. 2019;33(4):199–206.

18. Blasco M, et al. Complement Mediated Endothelial Damage in Thrombotic Microangiopathies. Front Med (Lausanne). 2022;9:811504.

19. Aiello S, et al. C5a and C5aR1 are key drivers of microvascular platelet aggregation in clinical entities spanning from aHUS to COVID-19. Blood Adv. 2022;6(3):866–881.

20. Yang J, et al. Insights Into Immunothrombosis: The Interplay Among Neutrophil Extracellular Trap, von Willebrand Factor, and ADAMTS13. Front Immunol. 2020;11:610696.

21. Trivioli G, Vaglio A. The rise of complement in ANCA-associated vasculitis: from marginal player to target of modern therapy. Clin Exp Immunol. 2020;202(3):403–406.

22. Afzali B, et al. The state of complement in COVID-19. Nat Rev Immunol. 2022;22(2):77– 84.

23. Yu J, et al. Complement dysregulation is associated with severe COVID-19 illness. Haematologica. 2022;107(5):1095–1105.

24. Fujimura Y, Holland LZ. COVID-19 microthrombosis: unusually large VWF multimers are a platform for activation of the alternative complement pathway under cytokine storm. Int J Hematol. 2022;115(4):457–469.

25. Skendros P, et al. Complement C3 inhibition in severe COVID-19 using compstatin AMY-101. Sci Adv. 2022;8(33):eabo2341.

26. De Leeuw E, et al. Efficacy and safety of the investigational complement C5 inhibitor zilucoplan in patients hospitalized with COVID-19: an open-label randomized controlled trial. Respir Res. 2022;23(1):202.

27. Almaani S, et al. ANCA-Associated Vasculitis: An Update. J Clin Med. 2021;10(7):1446.

28. Xiao H, et al. Alternative complement pathway in the pathogenesis of disease mediated by anti-neutrophil cytoplasmic autoantibodies. Am J Pathol. 2007;170(1):52–64.

29. Jayne DRW, et al. Avacopan for the Treatment of ANCA-Associated Vasculitis. N Engl J Med. 2021;384(7):599–609.

30. Michelfelder S, et al. The MFHR1 Fusion Protein Is a Novel Synthetic Multitarget Complement Inhibitor with Therapeutic Potential. J Am Soc Nephrol. 2018;29(4):1141–1153.

31. Noris M, et al. Dynamics of complement activation in aHUS and how to monitor eculizumab therapy. Blood. 2014;124(11):1715–1726.

32. Perico L, et al. SARS-CoV-2 Spike Protein 1 Activates Microvascular Endothelial Cells and Complement System Leading to Platelet Aggregation. Front Immunol. 2022;13:827146.

33. Kirsch T, et al. Endothelial-derived thrombospondin-1 promotes macrophage recruitment and apoptotic cell clearance. J Cell Mol Med. 2010;14(7):1922–1934.

34. Michailidou D, et al. Neutrophil extracellular trap formation in anti-neutrophil cytoplasmic antibody-associated and large-vessel vasculitis. Clin Immunol. 2023;249:109274.

35. Kettritz R, et al. Role of mitogen-activated protein kinases in activation of human neutrophils by antineutrophil cytoplasmic antibodies. J Am Soc Nephrol. 2001;12(1):37–46.

36. Pickering MC, et al. Uncontrolled C3 activation causes membranoproliferative glomerulonephritis in mice deficient in complement factor H. Nat Genet. 2002;31(4):424–428.

37. Kaur S, Roberts DD. Why do humans need thrombospondin-1? J Cell Commun Signal. [published online ahead of print: January 23, 2023]. 10.1007/s12079-023-00722-5.

38. Józsi M, et al. The C-terminus of complement factor H is essential for host cell protection. Mol Immunol. 2007;44(10):2697–2706.

39. Kim IS, et al. Dysregulated thrombospondin 1 and miRNA-29a-3p in severe COVID-19. Sci Rep. 2022;12(1):21227.

40. Cervia-Hasler C, et al. Persistent complement dysregulation with signs of thromboinflammation in active Long Covid. Science. 2024;383(6680):eadg7942.

41. Pimanda JE, et al. Role of thrombospondin-1 in control of von Willebrand factor multimer size in mice. J Biol Chem. 2004;279(20):21439–21448.

42. Turner NA, Moake J. Assembly and activation of alternative complement components on endothelial cell-anchored ultra-large von Willebrand factor links complement and hemostasis-thrombosis. PLoS One. 2013;8(3):e59372.

43. Noone DG, et al. Von Willebrand factor regulates complement on endothelial cells. Kidney Int. 2016;90(1):123–134.

44. Bettoni S, et al. Interaction between Multimeric von Willebrand Factor and Complement: A Fresh Look to the Pathophysiology of Microvascular Thrombosis. J Immunol. 2017;199(3):1021–1040.

45. Vanguri VK, et al. Thrombospondin-1 binds to polyhistidine with high affinity and specificity. Biochem J. 2000;347(Pt 2):469–473.

46. Resovi A, et al. Current understanding of the thrombospondin-1 interactome. Matrix Biol. 2014;37:83–91.

47. Burmeister A, et al. Impact of neutrophil extracellular traps on fluid properties, blood flow and complement activation. Front Immunol. 2022;13:1078891.

48. Tuck MK, et al. Standard operating procedures for serum and plasma collection: early detection research network consensus statement standard operating procedure integration working group. J Proteome Res. 2009;8(1):113–117.

49. Michelfelder S, et al. Moss-Produced, Glycosylation-Optimized Human Factor H for Therapeutic Application in Complement Disorders. J Am Soc Nephrol. 2017;28(5):1462– 1474.

50. Kadam AP, Sahu A. Identification of Complin, a novel complement inhibitor that targets complement proteins factor B and C2. J Immunol. 2010;184(12):7116–7124.

51. Yuan X, et al. Small-molecule factor D inhibitors selectively block the alternative pathway of complement in paroxysmal nocturnal hemoglobinuria and atypical hemolytic uremic syndrome. Haematologica. 2017;102(3):466–475.

52. Rogg M, et al. α-Parvin Defines a Specific Integrin Adhesome to Maintain the Glomerular Filtration Barrier. J Am Soc Nephrol. 2022;33(4):786–808.

