## Supplemental data for "Thrombospondin-1 inhibits alternative complement pathway activation in vasculitis synergistically to factor H"

### Methods:

#### Patient's description

Blood samples were collected from aHUS patients/relatives, PNH, ANCA-vasculitis patients and controls. Patients' details are described in supplemental table 1A-C

#### Supplemental Table 1

### A

#### aHUS

| | Age | Gender | Genetics | Autoantibodies | Creatinine (mg/dl) | Serum C3 (g/l) | Proteinuria (g/g creatinine) | Actual complement therapy |
|---|---|---|---|---|---|---|---|---|
| **aHUS1** | 34 | Male | *het. FH-* mutation R1215Q | No | 0,89 (0,6-1,2) | 1,1 (0,9-1,8) | 0,04 (<0,2) | No |
| **aHUS2** | 12 | Male | *homoz. CFHR1/3 deletion* | Anti-FH: 147U/ml (<10) | 2,46 (0,3 - 0,5) | 0,63 (0,9-1,8) | 3,03 (<0,2) | Ravulizumab[A]/ mycophenolate mofetil |
| **aHUS3** | 24 | Male | *het. FH-c-terminal deletion* | No | 2,64 (0,6 - 1,2) | <0,19 (0,9-1,8) | 0,24 (<0,2) | Ravulizumab[A] |

**Table S1A:** Summary of clinical data of aHUS patients/relatives: aHUS1 is a healthy man from an aHUS family carrying a previously described *FH* mutation. The *FH* mutation causes strong hemolytic activity when serum is incubated with sheep erythrocytes. Het. is heterozygous, homoz. is homozygous mutation, A - Serum samples used for in vitro assays were taken before the initiation of complement therapy

**B**

### PNH

| | Age | Gender | C59% negative erythrocytes (%) | Complement therapy |
|---|---|---|---|---|
| **PHN1** | 64 | Male | 99,5 | Iptacopan |
| **PHN2** | 77 | Female | *99,7* | Iptacopan |
| **PNH3** | 24 | Female | *47,9* | Pegcetacoplan |

**Table S1B:** Summary of clinical data of PNH patients

**C**

### ANCA-vasculitis/controls

| | Age | Gender | Histology<br>(crescent/sclerosis of total glomeruli) | Autoantibodies[A] | Creatinine[A]<br>(mg/dl) | Proteinuria[A]<br>(g/g creatinine) |
|---|---|---|---|---|---|---|
| **Patient diagnosis & number** | | | | | | |
| **Nephrektomy1<br>(Oncocytoma)** | 71 | Male | Normal<br>(0/0 of 15) | n.d. | 1,3<br>(0,6-1,2) | 0 |
| **Nephrektomy2<br>(Clear Cell Renal Cell Carcinoma)** | 68 | Male | Normal<br>(0/0 of 50) | n.d. | 4,3<br>(0,6-1,2) | n.d. |
| **Nephrektomy3<br>(Clear Cell Renal Cell Carcinoma)** | 64 | Male | Normal<br>(0/0 of 50) | n.d. | 2,1<br>(0,6-1,2) | 0,06<br>(<0,2) |
| **Nephrektomy4<br>(Clear Cell Renal Cell Carcinoma)** | 55 | Male | Normal<br>(0/0 of 50) | n.d. | 1,1<br>(0,6-1,2) | n.d. |
| **ANCA1** | 44 | Female | pauci-immun<br>(21/3 of 34) | Anti-PR3: >200U/ml<br>(<10) | 3,1<br>(0,5-1,0) | 1,18<br>(<0,2) |
| **ANCA2** | 21 | Female | pauci-immun<br>(5/3 of 13) | Anti-MPO: >200U/ml<br>(<20) | 1,2<br>(0,5-0,8) | 0,5<br>(<0,2) |
| **ANCA3** | 60 | Male | pauci-immun | Anti-PR3: >200U/ml | 2,8 | 2,94 |

| | | | | | | |
|---|---|---|---|---|---|---|
| | | | (7/3 of 12) | (<10) | (0,6-1,2) | (<0,2) |
| **ANCA4** | 63 | Male | pauci-immun<br>(1/0 of 6) | Anti-MPO: 120U/ml<br>(<20) | 5,8<br>(0,6-1,2) | 1,25<br>(<0,2) |
| **FSGS1**<br>**Transplant; FSGS relapse** | 15 | Female | FSGS<br>(0/7 of 50) | n.d. | 0,94<br>(0,5-1,0) | 6,52<br>(<0,2) |
| **FSGS2** | 65 | Male | FSGS<br>(0/11 of 15) | n.d. | 2,76<br>(0,6-1,2) | 1,50<br>(<0,2) |
| **FSGS3** | 54 | Male | FSGS<br>(0/4 of 9) | n.d. | 3,25<br>(0,6-1,2) | 2,49<br>(<0,2) |
| **FSGS4** | 19 | Female | FSGS<br>(0/50 of 50) | n.d. | 4,1<br>(0,5-1,0) | 7,15<br>(<0,2) |

**Table S1C:** Summary of clinical data of ANCA-vasculitis patients (ANCA), controls (nephrectomy and FSGS (focal segmental glomerular sclerosis). A - values at time of renal biopsy, n.d.: not determined

### ELISA confirming binding to complement proteins with platelet isolated TSP-1 (p-TSP-1)

Binding of TSP-1 to complement proteins was confirmed using untagged TSP-1 isolated from human platelets. Recombinant TSP-1 (TSP-1), platelet-isolated TSP-1 (p-TSP-1), Eculizumab, BSA or, where applicable, FH was coated overnight on Nunc maxisorb 96 well plates at 133 nM each in PBS. Unbound proteins were washed with PBS containing 0.05 % Tween20 and the wells blocked with PBS containing 2 % BSA for 1 h at room temperature. After washing, human FH, FB, C3 or C5 was added to the wells at 10 or 20 µg/ml and incubated for 2 h at room temperature. After washing, complement proteins were detected by incubating the samples for 2 h at room temperature with the following specific primary antibodies: goat anti-human FH (Complement Technology, A237), goat anti-human FB (Calbiochem, 341272), goat anti-human C3 (Complement Technology, A213), mouse anti-human C5 (Quidel, A306). After washing, the samples were incubated for 1 h at room temperature with rabbit anti-goat HRP

(Dako, P0449) or sheep anti-mouse HRP (GE-Healthcare, NXA931) secondary antibodies. Colorimetric detection of HRP antibodies was performed using TMB substrate. The reaction was stopped after 10 min and optical density at 450 nm measured. Cofactor and decay acceleration assay:

Cofactor activity of FH in the presence or absence of TSP-1 was assessed as described before (1). C3 cleavage products were visualized using coomassie staining after SDS-PAGE.

Decay acceleration activity of FH in the presence or absence of TSP-1 was measured by ELISA as previously described (2).

**Supplemental Figure 1: ELISA confirming BLI binding to complement proteins with platelet isolated TSP-1**

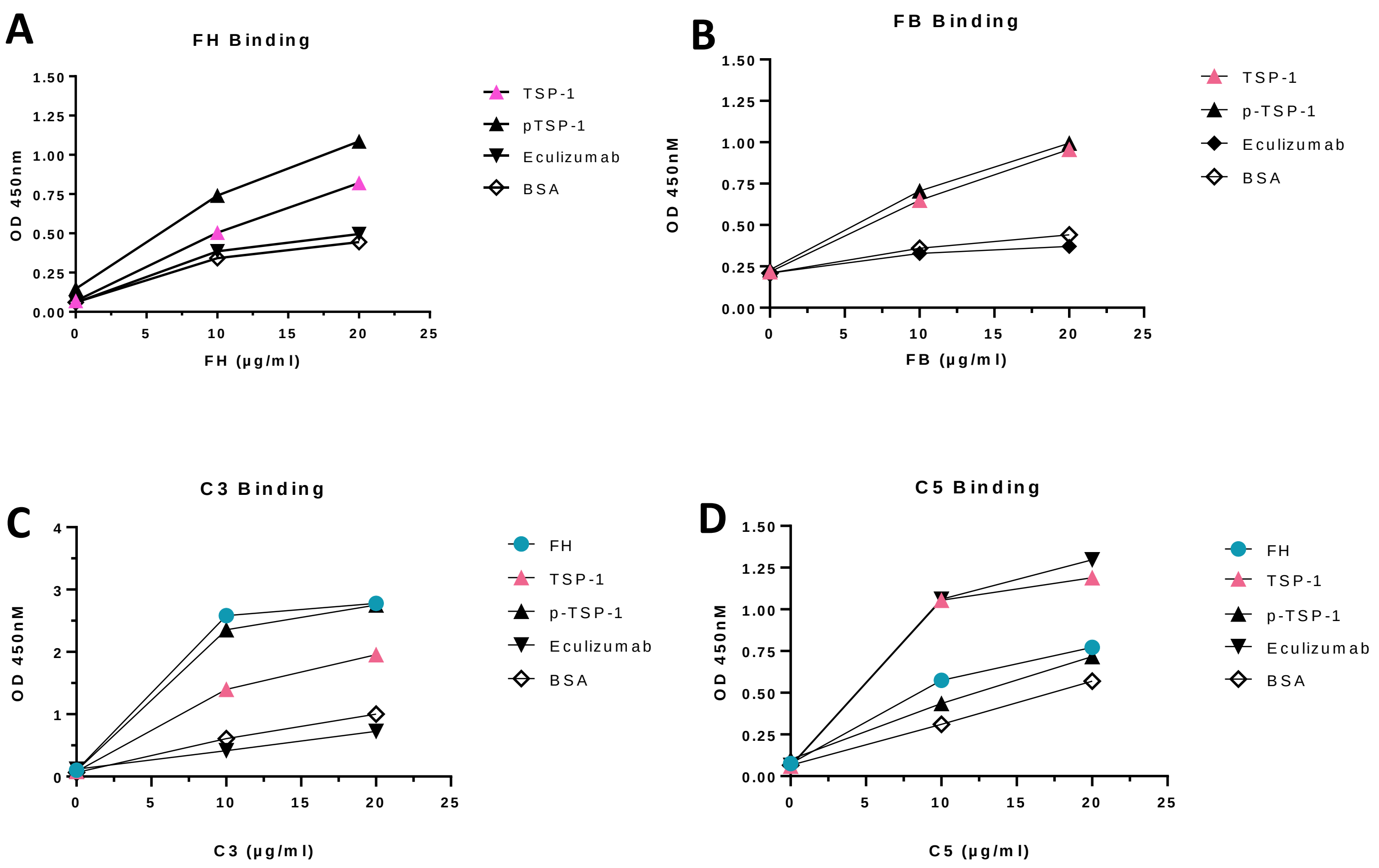


**Figure S1: Recombinant as well as platelet isolated TSP-1 binds to central alternative pathway proteins.** Recombinant TSP-1 (TSP-1), platelet-isolated TSP-1 (p-TSP-1), Eculizumab, BSA or FH were coated on microtiter plates at equimolar concentrations (133 nM). Complement proteins FH, FB, C3 or C5 were added to the wells at indicated concentrations and the amount of bound protein determined via ELISA. (A) Binding of TSP-1 and p-TSP-1 to complement protein FH. (B) Binding of TSP-1 and p-TSP-1 to complement protein FB. (C) Binding of TSP-1 and p-TSP-1 to complement protein C3. (D) Binding of TSP-1 and p-TSP-1 to complement protein C5. Results are shown as means.

**Supplemental Figure 2: Influence of TSP-1 on FH decay acceleration and cofactor activity:**

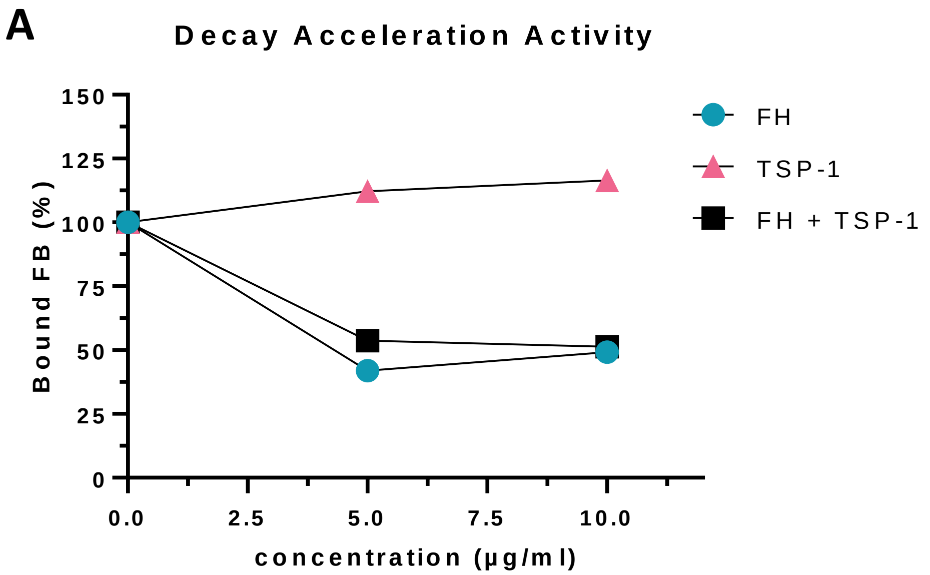


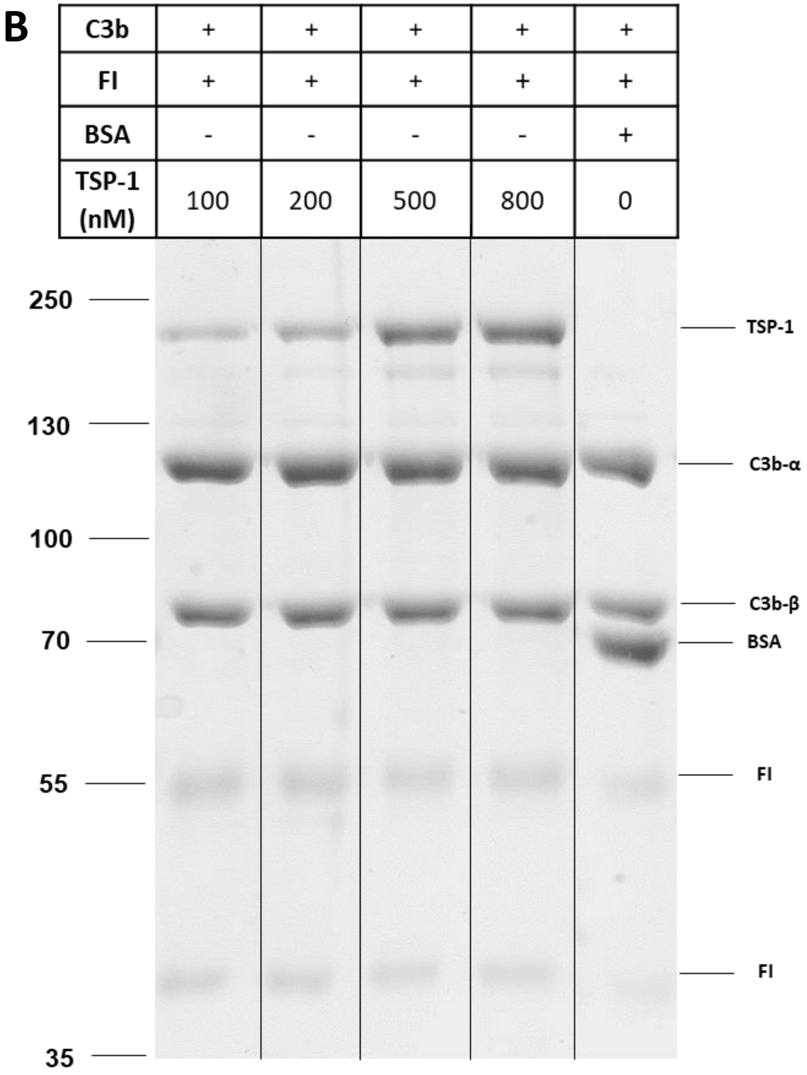


**Figure S2: TSP-1 does not influence decay acceleration activity of FH and has no intrinsic cofactor or decay acceleration activity.** (A) Decay acceleration activity of TSP-1. Microtiter wells were coated with C3b and C3 convertase was generated by adding FB and FD. Convertases were incubated with increasing concentrations of either FH alone, FH in the presence of 10 µg/ml TSP-1 or with increasing concentrations of TSP-1 alone. The amount of intact C3bBb molecules left on the surface detected with antibodies against FB. (B) Cofactor Activity of TSP-1. C3b and FI were incubated with increasing concentrations of TSP-1 or BSA. Cleavage of C3b was visualized on coomassie stained SDS gels. Results are shown as means.

**Supplemental references:**


1. Lambris JD, et al. A discontinuous factor H binding site in the third component of complement as delineated by synthetic peptides. *J Biol Chem*. 1988;263(24):12147–12150.

2. Michelfelder S, et al. Moss-Produced, Glycosylation-Optimized Human Factor H for Therapeutic Application in Complement Disorders. *J Am Soc Nephrol*. 2017;28(5):1462–1474.
